# Munc18 reprograms the intrinsic neuronal SNARE complex assembly pathway

**DOI:** 10.64898/2026.07.21.739636

**Authors:** Vicky Vishvakarma, Edwin R. Chapman

## Abstract

The SNARE proteins syntaxin/SNAP-25B (t-SNAREs) and synaptobrevin (v-SNARE) contain motifs that assemble into four-helix bundles to drive synaptic vesicle exocytosis; SNAP-25B contributes two helices, D1 and D2. The sequence in which these motifs interact remains unresolved. To address this, we used fluorescence anisotropy of SNARE motifs to conduct real-time order-of-addition experiments and found that the order in which components are mixed can determine whether on- or off-pathway complexes are formed. Beginning with soluble SNARE fragments alone, the first step in assembly is the binding of D1 to syntaxin, followed by the binding of synaptobrevin and D2, where the latter motif acts as a gatekeeper to control v-SNARE•t-SNARE interactions. We then examined the impact of two regulatory factors, Munc18 and the MUN domain of Munc13-1. Strikingly, in the presence of Munc18, all four isolated SNARE motifs must be present at the same time for assembly to occur, revealing a concerted mechanism, while the Munc13-1 fragment was without effect. We created C-SCORE, which reports assembly of the two SNARE motifs of SNAP-25B via FRET, and confirmed that in the presence of Munc18, SNARE assembly becomes concerted. Munc18 also disaggregated syntaxin, potentially contributing to its activation. Finally, our findings regarding SNARE motif folding were well-correlated with function using full-length SNAREs in *in vitro* lipid mixing assays. Hence, Munc18 acts as a molecular chaperone that directly promotes the concurrent assembly of SNARE proteins into functional fusion machines.

## Introduction

Intracellular membrane fusion reactions are catalyzed by soluble N-ethyl maleimide-sensitive factor attachment protein receptors (SNAREs) (1). At neuronal synapses, three SNAREs, syntaxin 1A (syntaxin), SNAP-25B, and synaptobrevin 2 (Syb), mediate the fusion of synaptic vesicles (SV) with the presynaptic plasma membrane, resulting in the release of neurotransmitters. Syb is present on the SV membrane and is called a v-SNARE (2), while syntaxin and SNAP-25B are present on the target membrane and are referred to as t-SNAREs (3). These proteins contain evolutionarily conserved sequences of sixty to seventy amino acids that consist of heptad repeats, termed SNARE motifs, that fold together to form highly stable four-helix bundles. The formation of *trans*-SNARE complexes, between v- and t-SNAREs, brings the two membranes into close proximity, yet how these interactions drive membrane fusion remains unclear. Insights can be gained by understanding the assembly pathway of neuronal SNAREs into complexes, but this issue is the subject of considerable debate.

Three distinct models for SNARE complex formation have been proposed. In the first model, assembly starts with a strong interaction between SNAP-25B and syntaxin, which forms a high-affinity binding site for Syb (4, 5). In the second model, assembly begins with a weak interaction between syntaxin and Syb; this low-affinity binary interaction is facilitated by the regulatory protein, Munc18, which brings the N-terminus of syntaxin and Syb into close proximity (6, 7). Then, SNAP-25B joins, completing the assembly (7). In the third model, SNAP-25B first interacts with Syb, and this binary complex then binds syntaxin (8). The first two models have been proposed based on *in vitro* biochemical experiments using purified proteins (4–7), while the third model was inferred from indirect experiments using permeabilized PC-12 cells (8).

Interestingly, while the parallel four-helix bundle that forms the core of the SNARE complex contains one helix each from syntaxin and Syb, SNAP-25B is unusual in that it contributes two helices, termed D1 and D2 (9). Regardless of the order of subunit assembly, SNARE complexes have been hypothesized to progressively zipper from their N- to C-termini to drive fusion (10, 11). However, these findings were derived from experiments using a partially assembled t-SNARE complex stabilized with parts of the SNARE motif of Syb. While this idea is appealing, direct support for progressive zippering is lacking. A complicating factor is the observation that SNARE motifs can also form complexes with different stoichiometries and subunit compositions (12–14). Namely, syntaxin and SNAP-25B can form t-SNARE complexes with a 2:1 stoichiometry, respectively (13, 15). In addition, the crystal structure of the SNARE motif of syntaxin (H3) and the D1 domain of SNAP-25B revealed a complex with a 2:2 stoichiometry (16). Even the question of whether a binary interaction between syntaxin and Syb occurs remains the subject of debate (17, 18). Moreover, it is unknown which, if any, of these non-canonical complexes serve as productive intermediates during SNARE complex assembly *in vivo*. Yet another layer of complexity was introduced when the two SNARE motifs of SNAP-25B, D1, and D2 were independently examined (19). D1 is partially structured at its N-terminus, while D2 is completely disordered (18, 20, 21). Contrasting reports conclude that either domain can support vesicular fusion (21, 22) or that both domains must be present at the same time to drive secretion (23). Furthermore, regulatory proteins are likely to differentially affect these distinct assembly pathways, an issue that has been even less thoroughly addressed. For example, the impact of Munc18 remains unclear, as both positive and negative roles in SNARE complex formation have been proposed (24–26). Together, these findings illustrate the challenges in determining the precise pathway that underlies SNARE complex assembly.

In the current study, we comprehensively evaluated the intrinsic ability of SNARE motifs to assemble into complexes and examined the impact of two key regulatory proteins, Munc18 and the MUN domain of Munc13-1 (Munc13-1_MUN_), as well as the regulatory Habc domain of syntaxin, on the assembly pathway(s). Previous studies were largely focused on equilibrium binding (10, 21) and non-equilibrium stopped-flow rapid mixing experiments to address the kinetics of binary and ternary interactions (18). Here, we took a different approach by conducting novel order-of-addition experiments of isolated SNARE motifs and regulatory factors and monitoring their interactions in real time. This approach is crucial because the order in which components are combined, and their relative concentrations, can affect the outcome. We extended this work to full-length SNAP-25B by creating a modified fluorescence reporter (C-SCORE) that monitors the assembly of the C-termini of its two SNARE motifs, into SNARE complexes, via Förster resonance energy transfer (FRET). Finally, we correlated our assembly pathway findings, which were based on soluble SNARE motifs, with function, using full-length proteins and lipid mixing assays. These experiments provide insights into the precise order of events that underlie functional SNARE complex assembly and, importantly, show that this intrinsic pathway is strikingly altered by Munc18 and the Habc domain of syntaxin.

## Results

### Cooperative assembly of Syb with syntaxin and the SNARE motifs of SNAP-25B

The purified cytoplasmic domains of SNARE proteins spontaneously assemble into SNARE complexes (**Fig. 1*A***) relatively slowly at µM concentrations (27, 28). To monitor these interactions in solution in real time, we followed the fluorescence anisotropy of labeled cytoplasmic domains, or isolated SNARE motifs, upon addition of binding partners and regulatory domains; binding to another protein increases the anisotropy of the labeled species by changing the rotational correlation time (29, 30). The main approach here is to conduct order-of-addition experiments, but we note that when examining, for example, the assembly of five components, there are >100 permutations regarding the order in which components can be mixed, so it was essential to judiciously select the most crucial combinations to study assembly.

**Fig. 1.**
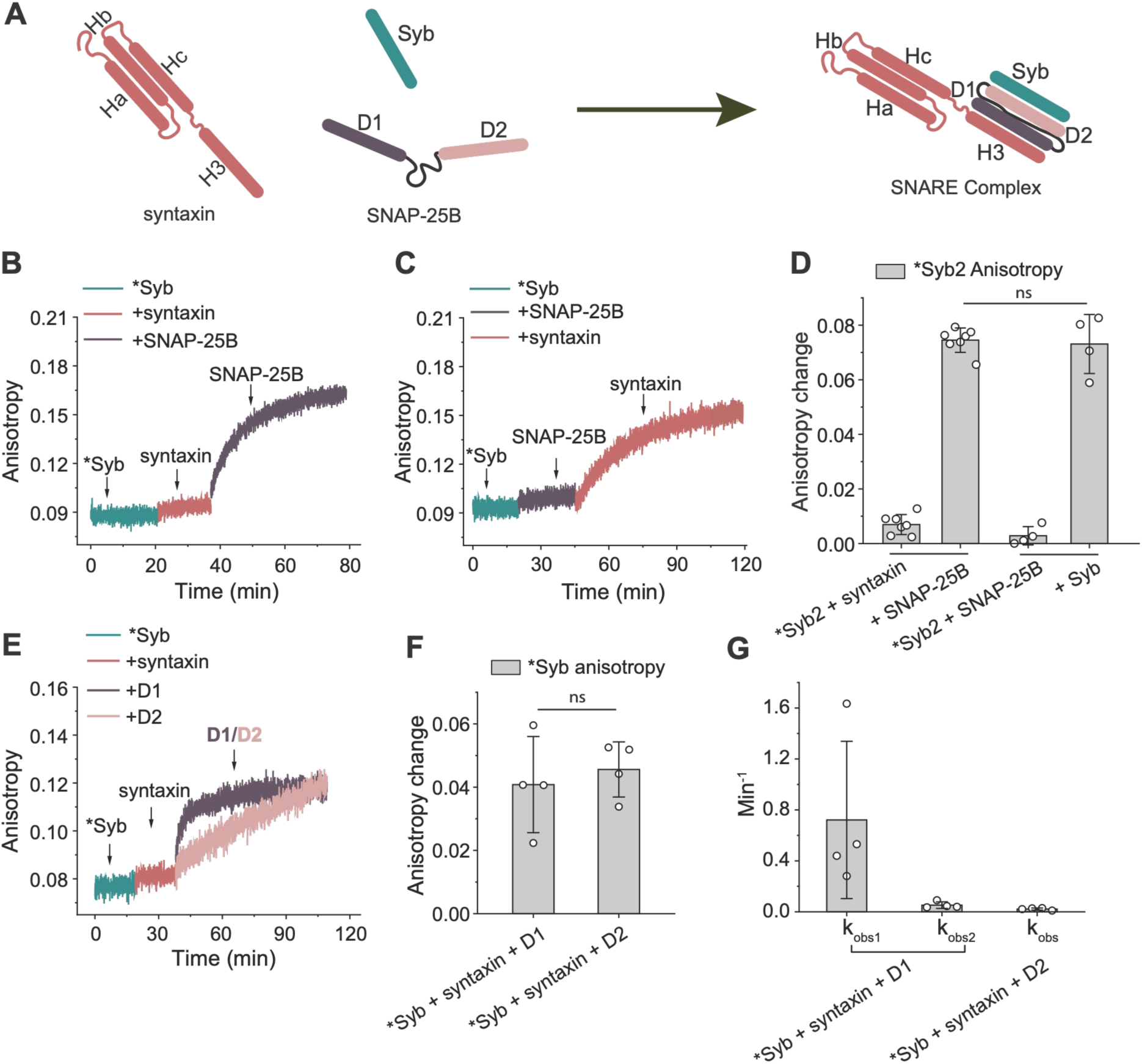
Syntaxin and both domains of SNAP-25B cooperate to bind synaptobrevin (Syb). (*A*) Illustration depicting assembly of the synaptic SNARE complex comprising syntaxin, SNAP-25B, and Syb. H3, D1, D2, and Syb are the SNARE motifs participating in four-helix bundle formation. (*B*) Fluorescence anisotropy trace of labeled (*) Syb (at amino acid 79) as a function of time, with sequential addition of syntaxin and SNAP-25B; n = 7. (*C*) Same as panel (*B*), but with sequential addition of SNAP-25B and syntaxin; n = 4. (*D*) Quantification of the change in Syb anisotropy, from the experiments shown in panels (*B*) and (*C*); The line segments at the bottom indicate the groupings of the same order-of-addition experiments. (*E*) Syb folds more rapidly with syntaxin-D1 than syntaxin-D2. Fluorescence anisotropy of labeled (*) Syb measured with sequential addition of syntaxin and D1 or D2 domain of SNAP-25B; the domain structure of SNAP-25B is illustrated panel (*A*). (*F*) Quantification of the fluorescence anisotropy changes of Syb with sequential addition of syntaxin and each SNARE motif (D1 (n = 4) and D2 (n = 4)) of SNAP-25B; values were obtained by averaging the trace between ∼ 60-70 minutes after adding D1 or D2. (*G*) Observed rate constants of Syb folding with syntaxin plus either the D1 or D2 fragment of SNAP-25B. The values were obtained by fitting the traces from panel (*E*) with single (D2) or double (D1) exponential functions. Error bars represent standard deviation. Statistical significance was determined by Welch’s unpaired t-test; ^#^*p* < 0.05, ^##^*p* < 0.01, ^###^*p* < 0.001, ^####^*p* < 0.0001, ns-not significant.

In the first series of experiments, we measured the fluorescence anisotropy of Syb, labeled at residue 79 near the C-terminus of its SNARE motif, with Alexa Fluor 488, and assayed for interactions with the cognate t-SNAREs, syntaxin and SNAP-25B. We observed a small increase in the fluorescence anisotropy of Syb upon addition of syntaxin (**Fig. 1*B***). However, when SNAP-25B was subsequently added, a robust time-dependent increase in anisotropy was observed (**Fig. 1*B*** and **1*D***). So, either all three components were required for assembly, or Syb strongly interacted with SNAP-25B under these conditions. To distinguish between these two possibilities, we repeated this experiment but reversed the order of addition of syntaxin and SNAP-25B (**Fig. 1*C*** and ***D***). Interestingly, a robust time-dependent increase in Syb anisotropy was still observed only when both the syntaxin and SNAP-25B were present. We note that the final anisotropy values, determined at the plateau phase, were same in both the experiments as shown in **Fig. 1*D***. These findings indicate that syntaxin and SNAP-25B cooperate to bind Syb during SNARE complex assembly.

SNAP-25B contributes two SNARE motifs, namely D1 and D2, to SNARE complexes. However, it was not clear if both motifs contributed to the increase in Syb anisotropy. To answer this question, we conducted the identical experiment as in **Fig. 1*B***, but replaced SNAP-25B with isolated D1 or D2. We observed that Syb folded faster onto syntaxin in the presence of the D1 domain of SNAP-25B as compared to the D2 domain (**Fig. 1*E, F, G***). These experiments demonstrated an inherent kinetic difference in the properties of the D1 and D2 domains; they were clearly non-equivalent (18, 21). The traces with D1 were readily fitted by a biexponential equation (revealing fast and slow kinetic components), whereas for D2, the data were well-fitted by a monoexponential function, revealing kinetics similar to the slow component exhibited by the D1 domain. The normalized amplitudes for fast and slow kinetic components of D1 were 0.54 ± 0.09 and 0.45 ± 0.09, respectively. These experiments indicate that either domain of SNAP-25B, D1 or D2, can cooperate with syntaxin to assemble with Syb into SNARE complexes, with D1 assembling before D2 into SNARE complexes.

There are conflicting reports regarding whether Syb directly binds syntaxin in binary reactions (17, 18). We addressed this issue by titrating syntaxin onto labeled Syb in the anisotropy assay. For completeness, we assessed this putative interaction by labeling Syb at two different positions, at either the aforementioned amino acid position 79, or toward the N-terminus at position 28. Titration of syntaxin resulted in an increase in Syb anisotropy, clearly demonstrating binding, albeit at relatively high [syntaxin], consistent with a low-affinity interaction in the binary complex lacking SNAP-25B (**Fig. S1)**. Notably, the anisotropy changes in Syb were markedly greater when labeled near its N-terminus as compared to its C-terminus. These results suggest that interactions in the binary complex are mediated by residues in the N-terminal region, consistent with N-to-C-terminal zippering.

### Distinct interactions of D1 and D2 with syntaxin

To delve deeper into the difference between the isolated D1 and D2 domains of SNAP-25B, we labeled them with a fluorescent probe at or near the N-terminal region of each individual domain: at residue 7 of the D1 domain, and at a cysteine added onto the N-terminus of the D2 domain. Upon addition of syntaxin, we observed a time-dependent increase in fluorescence anisotropy of labeled D1, demonstrating a direct interaction (**Fig. 2*A*** and ***D***). The robust anisotropy change indicates that D1 has a relatively strong affinity for syntaxin. In sharp contrast, the anisotropy of labeled D2 did not change upon addition of syntaxin; there was no measurable interaction under these conditions (**Fig. S2** and **Fig. 2*D***). Interestingly, D2 also did not exhibit detectable binding to either D1 or Syb (**Fig. 2*B*** and ***D***). The anisotropy value of D2 only changed in the presence of the syntaxin-D1 or the syntaxin-Syb pair (**Fig. 2*D*** and **Fig S2**). These findings highlight the utility of order-of-addition experiments and the idea that ternary interactions are necessary for the participation of D2 in SNARE complex assembly.

**Fig. 2.**
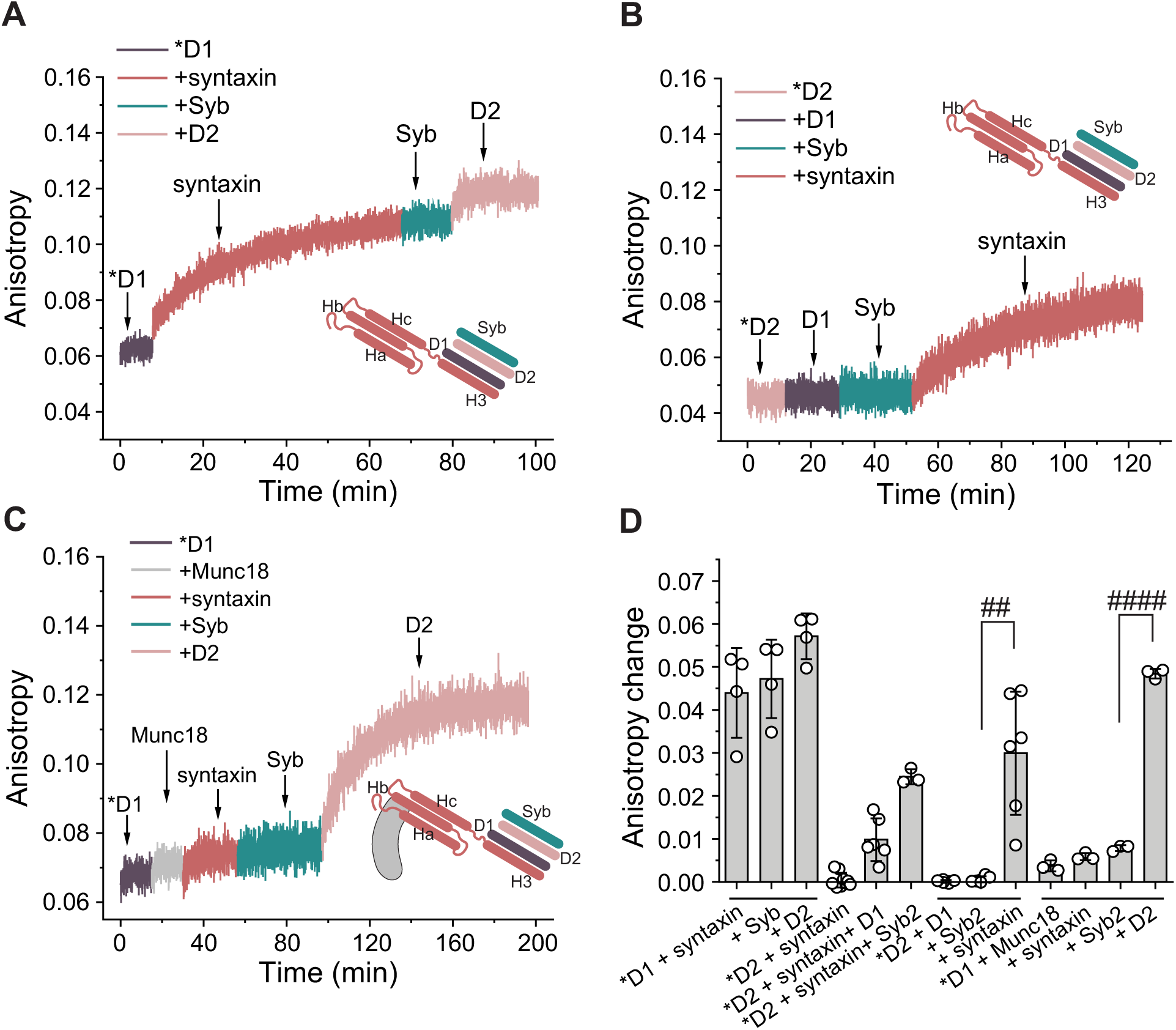
The D1 and D2 domains of SNAP-25B exhibit distinct binary and ternary interactions with syntaxin and Syb. (*A*) Representative trace of the fluorescence anisotropy of labeled (*) D1 measured with sequential addition of syntaxin, Syb, and the D2 domain of SNAP-25B; n = 4. (*B*) Similar experiment as in panel (*A*) but using the labeled (*) D2 domain of SNAP-25B with sequential addition of D1, Syb, and syntaxin; n = 6. Note: *D2 does not bind syntaxin, so the increase in anisotropy in panel (*B*) is due to the interaction of *D2 with D1, Syb, and syntaxin complexes (**Fig. S*2*)**. (*C*) Similar experiment as in panel (*A*) with the sequential addition of Munc18 and syntaxin, followed by the addition of Syb and D2; n = 3. (*D*) Quantification of the fluorescence anisotropy changes of the labeled D1 or D2 domains of SNAP-25B from panels (*A*, *B*, *C*), and **Fig. S2**. Error bars represent the standard deviation. The line segments at the bottom indicate the grouping of the same order-of-addition experiments. Note: in panel (*D*), the data sets lacking a line segment have different n values because data from additional experiments were pooled. Statistical significance was determined by Welch’s unpaired t-test; ^#^*p* < 0.05, ^##^*p* < 0.01, ^###^*p* < 0.001, ^####^*p* < 0.0001, ns-not significant.

Overall, these results suggest that a binary interaction exists between D1 and syntaxin, whereas D2 shows ternary and quaternary interactions with other SNARE motifs during SNARE complex assembly, with no significant binary interactions. Moreover, syntaxin appears to be the template for the SNARE motifs of SNAP-25B, as without syntaxin, individual D1 and D2 domains neither interact with each other nor with Syb. We note that Syb did not bind with the syntaxin-D1 complex in **Fig. 2*A***, which might seem to contradict **Fig. 1*E***. However, these observations can be explained by the formation of a syntaxin-D1 dead end complex in **Fig. 2*A***, as low concentrations of labeled D1 were saturated with syntaxin, likely forming 2:2 complexes (16) that are refractory to the binding of Syb. In contrast, the experiment in **Fig. 1*E*** began with a low concentration of labeled Syb, which binds to syntaxin and D1 as they are serially added; presumably, this occurs rapidly, before syntaxin and D1 can assemble into 2:2 off-pathway complexes. These findings, again, underscore the importance of the order of addition during SNARE complex assembly.

### Munc18 switches SNARE complex assembly from a sequential to a concerted pathway

In the next series of experiments, we studied the potential impact of the key regulatory proteins, Munc18 and the MUN domain of Munc13-1 (Munc13-1_MUN_), on the sequence of events that underlie SNARE complex assembly. We performed an experiment analogous to **Fig. 2*A***, but in the presence of these two regulatory proteins. Surprisingly, Munc18 blocked the robust interaction between labeled D1 and syntaxin (**Fig. 2*C*** and ***D***); it also blocked ternary interactions between D1, syntaxin, and Syb. Remarkably, Munc18 converts the assembly pathway from the sequential order of syntaxin-D1, followed by entry of Syb and D2 into SNARE complexes, into a concerted mechanism in which all four SNARE motifs must be present at the same time for complex assembly to occur. Hence, a single regulatory protein can reprogram the SNARE assembly pathway. In contrast, Munc13-1_MUN_ had no effect on D1-syntaxin interactions and did not affect SNARE complex formation under our experimental conditions (**Fig S3**).

### Role of D2 during SNARE complex assembly in the absence of regulatory proteins

Results from **Fig. 2** highlight the central role of syntaxin in SNARE complex assembly. To further investigate whether individual SNARE motifs assemble sequentially or simultaneously onto syntaxin, we measured the fluorescence anisotropy of its isolated SNARE motif, H3, labeled at its C-terminus via an added cysteine residue, over time with the sequential addition of other SNARE motifs. We observed a larger change in fluorescence anisotropy of H3 with SNAP-25B than with Syb, as shown in **Fig. 3*B*** and **Fig S4*A***, suggesting that the former binds more tightly. A caveat of this interpretation is that SNAP-25B is larger than Syb and so will affect the anisotropy, upon binding, to a greater extent. However, the literature supports the conclusion that the cytoplasmic domain of syntaxin binds more tightly to SNAP-25B than Syb (31), and these observations are likely to apply to the isolated H3 domain. Notably, when we measured the anisotropy of fluorescently labeled Syb with unlabeled H3, we observed a higher change in Syb anisotropy as compared to adding Syb to labeled H3 (**Fig S4**). Here, it is important to note that the labeled species is always at a low concentration in these experiments (200-300 nM), whereas the unlabeled species is always present at higher concentrations (∼10 µM, unless otherwise indicated). So, the difference in anisotropy values for labeled H3 versus Syb, upon addition of the other species, is consistent with the idea that at relatively high concentrations, H3 forms dimers or higher-order oligomers (16) that, due to their greater mass, have a much larger impact on the anisotropy of labeled Syb; it is also possible that H3 oligomers bind more tightly to Syb. To confirm whether H3 exists as a multimer at higher concentrations in our experiments, we measured the fluorescence anisotropy of 300 nM labeled H3 in the presence of 13 μM unlabeled H3. As shown in **Fig S5**, we observed a significant increase in fluorescence anisotropy, showing that H3 does, in fact, oligomerize (16) at the µM concentrations used in some of our experiments (e.g., **Fig S4*B***). Therefore, to study the interaction of H3 with either D1 or D2, we used labeled H3 at a low concentration, to minimize multimerization, and added relatively high concentrations of each of the SNAP-25B SNARE motifs. This approach revealed major differences between D1 and D2. We observed a strong increase in the anisotropy of H3 in the presence of D1 (**Fig. 3*A*** and ***C***). However, the addition of Syb to the H3-D1 complex yielded only a small increase in anisotropy (0.045 ± 0.004 to 0.050 ± 0.003), and subsequent addition of D2 was without effect. The small change with Syb could be due to weak direct interactions with *H3, or due to interactions with a small sub-population of H3-D1 complexes that form on-pathway complexes. However, the major finding here is that H3-D1 interactions tend to result in off-pathway/dead-end complexes. Indeed, as noted in above, H3 and D1 can form a kinetically stable complex with a 2:2 stoichiometry (16). So, we allowed D2 to interact with the H3-D1 complex for 12 hours and observed a significant decrease in H3 anisotropy (**Fig. 3*C***). These findings are consistent with the slow displacement of labeled H3 from the putative 2:2 H3-D1 off-pathway complex. Regardless, the majority of *H3-D1 complexes are off-pathway products.

**Fig. 3.**
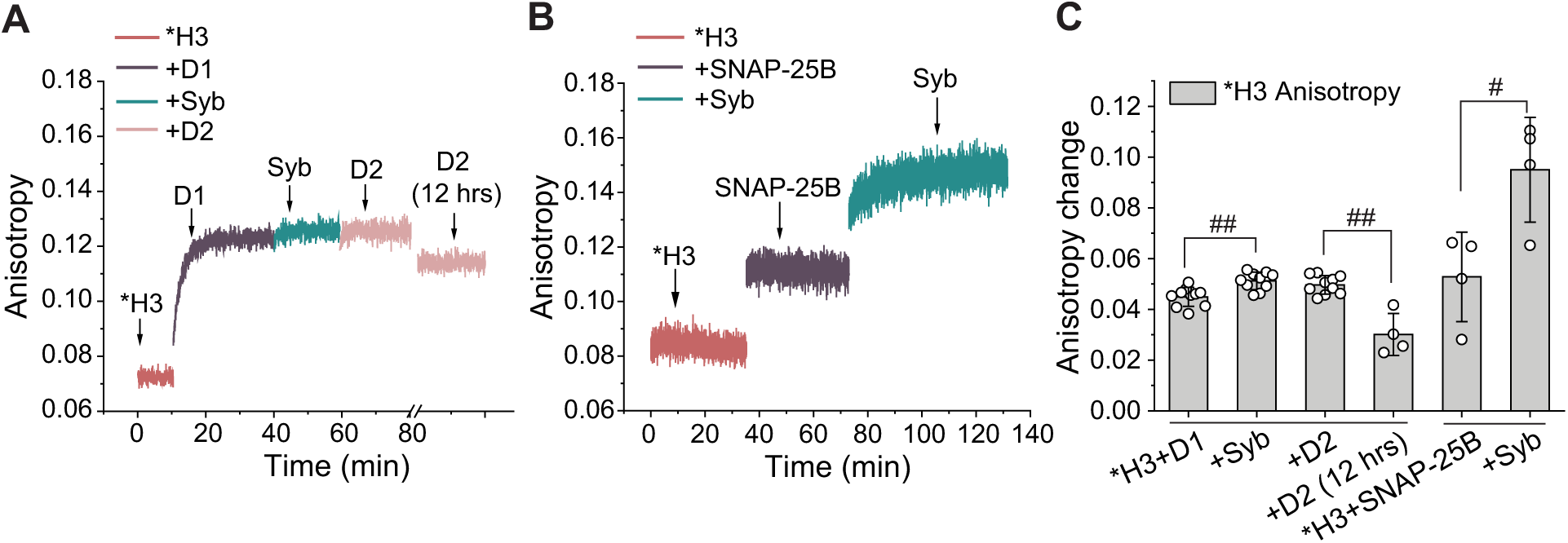
The SNARE motif of syntaxin (H3) and the D1 domain of SNAP-25B form a dead-end/off-pathway complex. (*A*) Fluorescence anisotropy of labeled (*) H3 measured with sequential addition of the D1, Syb, and the D2; n = 10. (*B*) Fluorescence anisotropy of labeled (*) H3 measured with sequential addition of SNAP-25B and Syb; n = 4. (*C*) Quantification of the fluorescence anisotropy changes of H3 from panel (*A*) and (*B*). The anisotropy of *H3 decreased 12 hours after addition of 10 µM D2 in panel (*A*) (n = 4 for this condition). The error bars represent the standard deviation. The line segments at the bottom indicate the grouping of the same order-of-addition experiments. Note: the +D2 (12 hrs) condition in panel (*C*) has only four data points because not all trials were completed at the 12 hrs time point. Statistical significance was determined by Welch’s unpaired t-test; ^#^*p* < 0.05, ^##^*p* < 0.01, ^###^*p* < 0.001, ^####^*p* < 0.0001, ns-not significant.

Syb exhibited a strong interaction with the H3-SNAP-25B complex, as shown in **Fig 3*B***, but had little, if any, interaction with syntaxin-D1 (**Fig. 2*A***) or H3-D1 (**Fig. 3*A***) complexes. These findings motivated us to further investigate the role of the D2 domain of SNAP-25B in the regulation of SNARE complex assembly. To address the role of the D2 domain, we designed three truncated constructs of SNAP-25B comprising residues 1-137, 1-180, and 1-197; the full-length protein is 206 residues. We refer to these fragments as D1-Linker, SNAP-25B_1-180_, and SNAP-25B_1-197_, respectively. SNAP-25B_1-180_ and SNAP-25B_1-197_ are the botulinum neurotoxin E (BoNT/E) (32) and botulinum neurotoxin A (BoNT/A) (33) cleavage products, respectively; D1-Linker comprises the D1 domain and the linker that connects D1 and D2. We observed an appreciable increase in the anisotropy of H3 upon addition of each of the three truncated forms of SNAP-25B (**Fig. 4**), demonstrating binding. We then added Syb and observed a slight increase (0.035 ± 0.005 to 0.043 ± 0.002) in the anisotropy of the H3-D1-Linker complex (**Fig. 4*A*** and ***D***). In contrast, a greater increase in anisotropy was observed for Syb with H3-SNAP-25B_1-180_ (0.036 ± 0.001 to 0.051 ± 0.003) (**Fig. 4*B*** and ***D***) and H3-SNAP-25B_1-197_ (0.016 ± 0.003 to 0.051 ± 0.009) (**Fig. 4*C*** and ***D***) complexes. So, extension of the SNAP-25B sequence to include portions of the D2 domain can facilitate the robust entry of Syb into the complex. Hence, D2 appears to act as a “gate-keeper” that regulates v-/t-SNARE pairing.

**Fig. 4.**
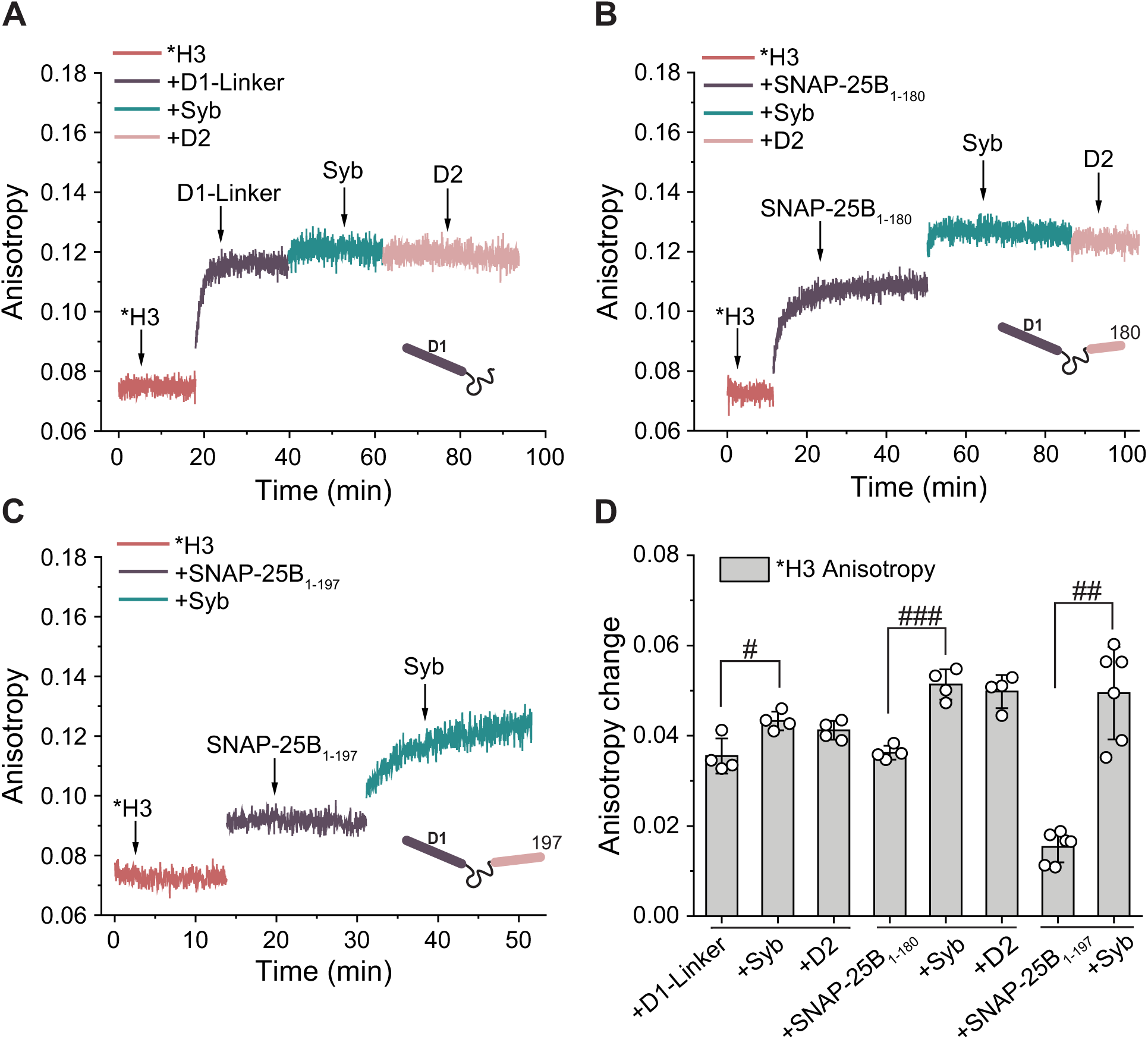
The C-terminal SNARE motif of SNAP-25B acts as a gatekeeper that controls assembly with Syb with H3. Fluorescence anisotropy of labeled (*) H3 measured with sequential addition of different fragments of SNAP-25B: (*A*) D1-Linker; n = 4, (*B*) SNAP-25B_1-180_; n = 4, and (*C*) SNAP-25B_1-197_; n = 6; Syb was subsequently added to each sample, and in panels (*A, B*); D2 was also added at the end of the experiment. (*D*) Quantification of the change in fluorescence anisotropy of labeled H3 calculated from the respective panels (*A*), (*B*), and (*C*). Error bars indicate the standard deviation. The line segments at the bottom indicate the grouping of the same order-of-addition experiments. Statistical significance was determined by Welch’s unpaired t-test; ^#^*p* < 0.05, ^##^*p* < 0.01, ^###^*p* < 0.001, ^####^*p* < 0.0001, ns-not significant.

We note that the interaction of SNAP-25B_1-197_ with H3 resulted in a smaller change in H3 fluorescence anisotropy as compared to both full-length SNAP-25B and SNAP-25B_1-180_ **(Fig 3*B*, *C***; **Fig. 4*B*** and ***D***). The reasons for this observation are unclear and bear further study. Regardless, these data reveal that extending D1 to contain portions of D2, in SNAP-25B_1-180_ and SNAP-25B_1-197_, aids the entry of Syb. While extension of SNAP-25B to include more of the D2 domain facilitated v-SNARE assembly into the complex, terminal addition of isolated D2 in **Fig. 4*A-B***, did not result in any change in the H3 anisotropy. This could be due to partial occupancy of the D2 binding site by SNAP-25B_1-180_ (**Fig. 4*B***), but this does not apply to the experiment using D1-linker (**Fig. 4*A***); perhaps D2 can enter on longer time scales, or this represents an off-pathway. This will be addressed in future studies by using higher concentrations of D2 and examining interactions over a longer time period.

### Munc18 works in conjunction with Habc to prevent off-pathway complex formation

Comparing **Fig. 1*B*** and **Fig S4*B*,** we found that the Habc domain of syntaxin inhibits binary interactions between the H3 domain and Syb, perhaps by forming a closed version of syntaxin (16). Additionally, the Habc domain also interacts with the regulatory protein Munc18 and the MUN domain of Munc13-1 (Munc13-1_MUN_), and these interactions are thought to play a key role in activating syntaxin during SNARE complex assembly *in vitro* (34, 35) and *in vivo* (36). Therefore, we explored the effects of Habc, Munc13-1_MUN_, and Munc18 on the SNARE complex assembly pathway and on the H3-D1 off-pathway. We first measured the fluorescence anisotropy of labeled H3 upon sequential addition of these regulatory proteins/fragments. H3 anisotropy increased in the presence of the Habc domain, confirming this well-characterized intra-domain interaction in syntaxin (24). The H3-Habc complex exhibited a strong interaction with D1, and Syb was now also able to bind to this complex when sequentially added (**Fig S6*A*** and ***B***). These findings demonstrate a positive role played by Habc by allowing Syb to bind to the H3-D1 complexes. Subsequent addition of D2 had little effect, because it binds on very slow time scales (**Fig. 3**), or because – under these conditions – dead-end complexes, with unknown stoichiometry, were formed. Our findings regarding Habc contrast with earlier work showing that removal of the Habc domain accelerates liposome fusion *in vitro* (37, 38), (4). However, in those experiments, syntaxin and SNAP-25B were co-expressed, so off-pathway complex formation was unlikely to occur.

Next, we investigated the effect of the MUN domain of Munc13-1 (35, 39), which interacts with both the SNARE motif of syntaxin (34) and the linker region of SNAP-25B (40). If Munc13-1_MUN_ blocks off-pathway H3-D1 complex formation, we should observe an increase in the H3 anisotropy upon addition of Syb. We observed a slight increase in fluorescence anisotropy of labeled H3 in the presence of the Munc13-1_MUN_ domain, suggesting a weak interaction; we then observed a significant increase in anisotropy when D1-Linker was added. However, we did not observe any increase in anisotropy when Syb and D2 were subsequently added (**Fig S6*C*** and ***D***), indicating that Munc13-1_MUN_ did not affect the formation of the off-pathway complex between H3 and D1. Either the linker region of SNAP-25B is not involved in the formation of this complex, or the interaction between the linker region and Munc13-1_MUN_ is too weak to inhibit the off-pathway under our experimental conditions.

Turning to Munc18, we observed a slight increase in H3 anisotropy, again suggesting a weak interaction, consistent with previous reports (41). Then, sequential addition of D1-Linker to the H3-Munc18 complex caused a significant increase in the fluorescence anisotropy of H3 (**Fig S6*E*** and ***F*)**. However, no further changes were observed upon sequential addition of Syb and D2, suggesting that Munc18 also did not affect the formation of the off-pathway complex between isolated SNARE motifs, H3, and D1-Linker.

The interactions between Munc13-1_MUN_ and Munc18 with syntaxin are influenced by the Habc domain of syntaxin (34, 35). We therefore monitored the assembly of SNARE motifs into complexes in the presence of all three of these regulatory factors. We again found a slight increase in labeled H3 anisotropy upon addition of Munc13-1_MUN_, and a significant increase with further addition of the Habc domain, suggesting unperturbed H3-Habc interactions in the presence of Munc13-1_MUN_. Subsequent addition of Munc18 did not change the H3 anisotropy (**Fig. 5*A*** and ***C***), despite the fact that it likely forms tight complexes with ‘closed’ syntaxin, which is presumably formed when Habc binds H3 (25). The reasons for the lack of an increase in anisotropy are complex and might include the finding that Munc18 greatly reduced the diffusion coefficient of the Habc-H3 complex via disaggregation (**Fig. 6*A*** and ***B***), potentially off-setting anisotropy increases in H3.

**Fig. 5.**
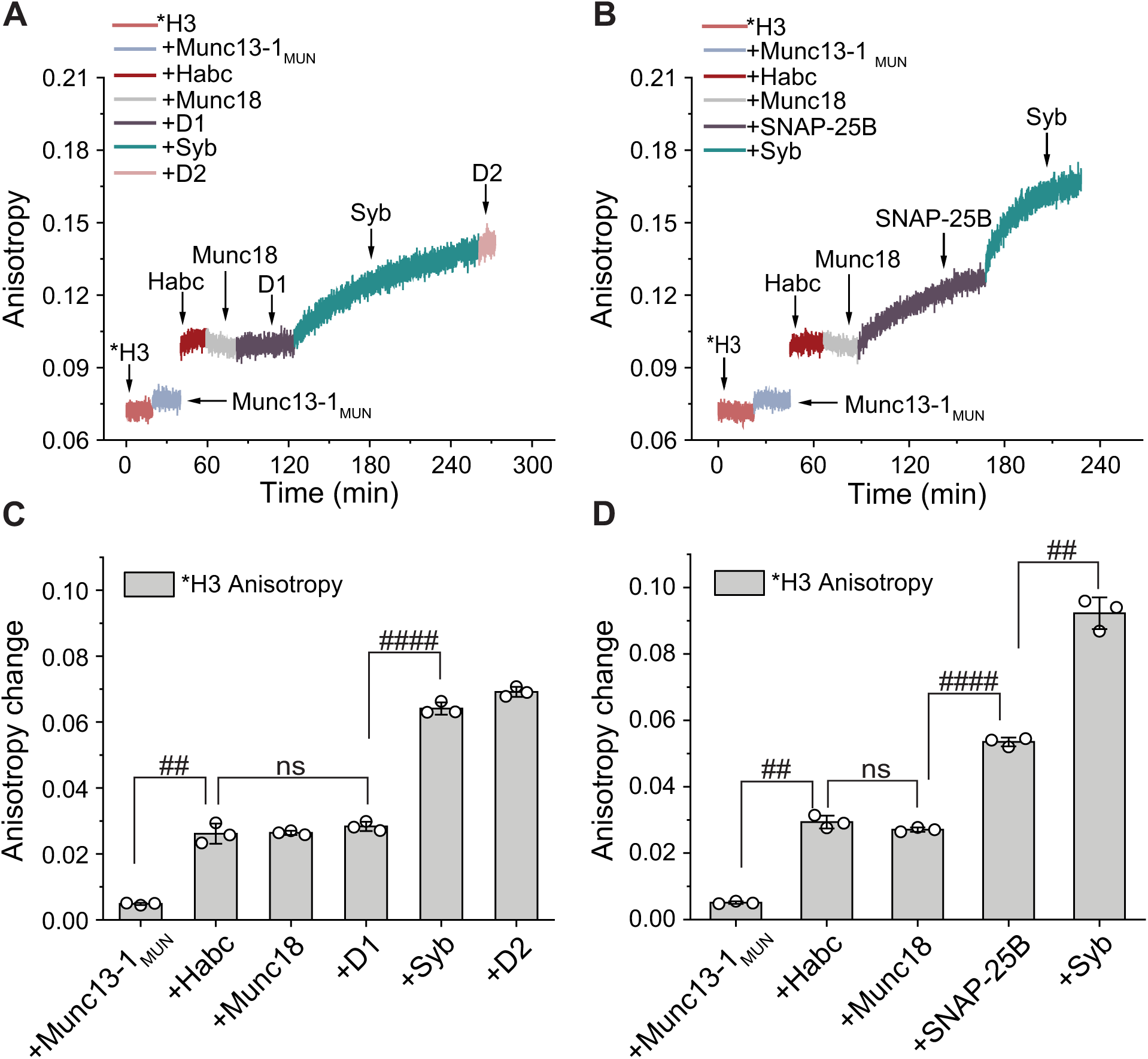
Munc18 and the Habc domain of syntaxin reprogram the SNARE complex assembly pathway. (*A*) Fluorescence anisotropy trace of labeled (*) H3 measured with sequential addition of the Munc13-1_MUN_, the syntaxin Habc domain, full-length Munc18, D1, Syb, and D2; n = 3. (*B*) Same experiment as in panel (*A*), but using full-length SNAP-25B instead of D1 or D2; n = 3. (*C*, *D*) Quantification of the change in fluorescence anisotropy of H3 corresponding to the experiments shown in panels (*A*) and (*B*), respectively. Error bars represent the standard deviation. Statistical significance was determined by Welch’s unpaired t-test; ^#^*p* < 0.05, ^##^*p* < 0.01, ^###^*p* < 0.001, ^####^*p* < 0.0001, ns-not significant.

**Fig. 6.**
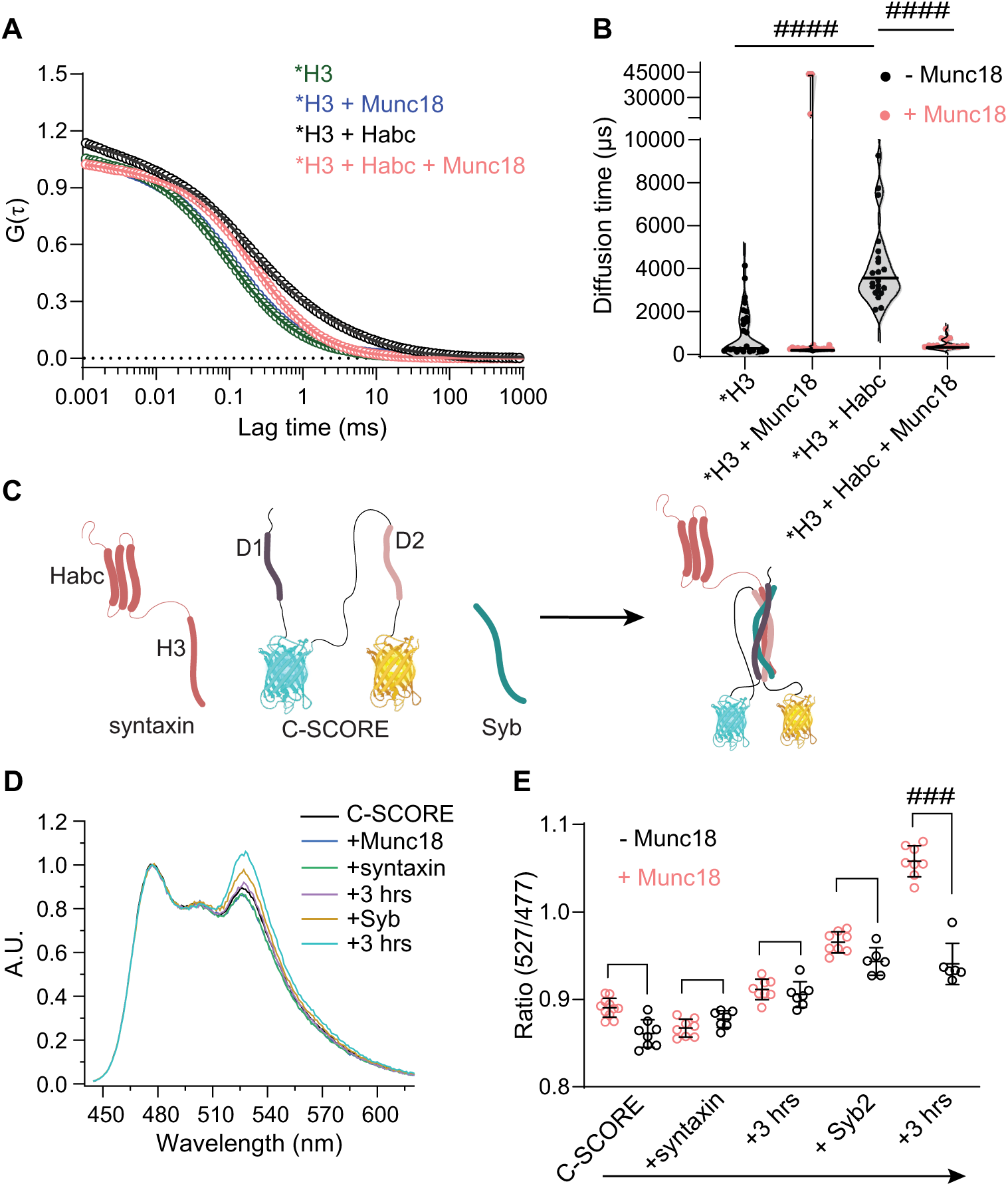
Munc18 promotes concerted SNARE complex formation; correlation with the inhibition of syntaxin oligomerization. (*A*) Representative FCS trace of labeled H3 with and without Habc and/or Munc18. (*B*) Bar graph showing the diffusion time estimated from panel (*A*). Statistical significance was determined by Mann-Whitney U test; ^#^p < 0.05, ^##^p < 0.01, ^###^p < 0.001, ^####^p < 0.0001, ns-not significant. (*C*) Illustration showing SNARE complex formation by syntaxin, C-SCORE, and Syb; the fluorescent proteins were created using BioRender.com. (*D*) Fluorescence spectra of C-SCORE during sequential addition experiments in which Munc18, syntaxin, and Syb were added in series. After each addition, the spectra were immediately collected and then re-measured after the indicated 3 hrs incubation period. Spectra were normalized to the cyan peak (477 nm); increases in the venus peak (527 nm), due to FRET, report assembly of D1 and D2 into the complex. (*E*) The FRET ratios (527/477) from panel (*D*) were calculated and plotted; each data point represents an independent measurement, and error bars represent the standard deviation. Statistical significance was determined by Mann-Whitney U test; ^#^*p* < 0.05, ^##^*p* < 0.01, ^###^*p* < 0.001, ^####^*p* < 0.0001, ns-not significant.

When Munc18, Habc, and Munc13-1_MUN_ were present, addition of the isolated D1 domain did not change the H3 anisotropy, consistent with the inability of D1 to bind syntaxin in the presence of Munc18 in **Fig. 2*C***. However, we observed a slow and significant increase in H3 anisotropy upon subsequent addition of Syb, showing that this SNARE motif was able to enter the complex, perhaps in conjunction with D1, in the presence of all three regulatory factors. We then conducted an analogous experiment, but using full-length SNAP-25B instead of the isolated D1 and D2 domains. SNAP-25B caused a slow and significant increase in H3 anisotropy when Munc18, Habc, and Munc13-1_MUN_ were present (**Fig. 5*B*** and ***D***). These results indicate that H3-SNAP-25B interactions are stronger than H3-D1 interactions in the presence of all three regulatory factors under these conditions. We note that H3-SNAP-25B interactions are not blocked by Munc18 (**Fig S7)**, and Munc13-1_MUN_ did not inhibit the interaction between syntaxin and D1 (**Fig S3)**. Moreover, when tested individually in the absence of the Habc domain, neither Munc18 nor Munc13-1_MUN_ inhibited H3-D1-Linker interactions (**Fig S6)**. Together, these findings reveal that dead-end H3-D1 interactions were inhibited due to interactions between Munc18 and the Habc domain of syntaxin. These findings further highlight the relevance of the Munc18-Habc interactions during SNARE complex assembly and the importance of linking the D1 and D2 domains of SNAP-25B together. These observations also shed further light on how Munc18 and Habc might convert the SNARE complex assembly pathway from a sequential to a concerted mechanism, as discussed above (**Fig. 2*C*** and ***D***).

### Munc18 prevents the formation of off-pathway t-SNARE complexes and promotes the concerted folding of SNAREs

Our real-time folding results suggest that Munc18 controls SNARE complex assembly primarily by acting on the Habc domain of syntaxin. In the literature, Munc18 has been shown to bind to the closed conformation of isolated syntaxin (42) as well as with the open form that is fully assembled into SNARE complexes (43). Given these multiple modes of binding, it remains unresolved how Munc18 orchestrates SNARE complex assembly. To investigate the underlying mechanism, we first examined the effect of Munc18 on syntaxin using an in-house custom-built fluorescence correlation spectrometer (FCS) (44). We monitored the interaction of fluorescently labeled H3 with Habc, in the presence and absence of Munc18. Normalized representative FCS traces are shown in **Fig. 6*A***. We analyzed the data following a two-component 3D-diffusion model (equation 1, see Methods for details) and observed that H3, alone shows two distinct populations with different diffusion times: a short timescale component *τ_D1_*, and a long timescale component, *τ_D2_*, with *g1* and *g2* being the amplitudes, respectively. We designate *τ_D1_* as diffusion time of monomeric/small-oligomeric H3 while *τ_D2_* > 1000 μs represents larger H3 oligomers. Violin plots of the long timescale diffusion time (*τ_D2_*) obtained from each experimental trial are shown in **Fig. 6*B***. The fractional population of the larger H3 oligomers [*g_2_*/(*g_1_*+*g_2_*)], estimated from trials showing *τ_D2_* > 1000 μs, was 0.1 ± 0.06 (mean ± SD), see **Table S1**. It is apparent that in most experimental trials, *τ_D2_* was greatly reduced by the addition of Munc18, indicating that Munc18 dissolved the large H3 oligomers. In the presence of Habc, H3 forms even bigger oligomers (fractional population ∼ 0.26 ± 0.02, mean ± SD) that, interestingly, are also dissolved by Munc18, as the average diffusion time was reduced ∼ 9-fold by this regulatory protein. These results suggest that Munc18 inhibits the oligomerization of syntaxin by interacting with its SNARE motif and Habc domain, to form an intermediate that is receptive for the concurrent binding of syntaxin and Syb. Our FCS results are consistent with a recent report in which Munc18 has been reported to modulate the liquid-liquid phase separation of syntaxin (45).

The anisotropy experiments above are appropriate for probing intrinsic SNARE interactions, but were limited to using soluble SNARE motifs/fragments. Moreover, assembly of unlabeled isolated D2 domain into on/off pathway complexes was difficult to observe in some of our experiments, due to weak interactions with other isolated SNARE motifs, or because the kinetics of binding were too slow to detect during the observation window. To better understand the folding of full-length SNAP-25B, in which D1 and D2 are tethered together, we engineered it to harbor a FRET (Förster resonance energy transfer) pair that reports its assembly into SNARE complexes. Namely, we introduced a cyan fluorescent protein (CFP) at the C-terminus of the D1 domain and a yellow fluorescent protein (Venus) at the C-terminus of the D2 domain (**Fig. 6*C***) in the intact protein. We designate this reporter as C-SCORE, an abbreviation for <u>C</u>-terminally tagged <u>S</u>NARE <u>CO</u>mplex <u>RE</u>porter, to distinguish it from previous analogous constructs (46–48) that did not report efficient FRET in our hands.

As shown in **Fig. 6*D***, we monitored the fluorescence spectrum of C-SCORE during sequential addition experiments; these are the first experiments, in the current study, where the action of Munc18 was studied using the complete cytoplasmic domains of all three SNARE proteins. We first added syntaxin, and then waited for three hours, followed by the addition of Syb, followed by another three-hour incubation period. At each step, we calculated the fluorescence intensity ratio (527/477) as a measure of FRET between CFP and Venus. We observed a higher FRET ratio in the presence of Munc18 when all four SNARE motifs were present concurrently; that is, SNAP-25B assembled with syntaxin only after the addition of the third SNARE, in this case Syb. Notably, in the absence of Munc18, addition of Syb had little effect on the FRET signal under otherwise identical conditions. These findings can be explained by the formation of a syntaxin•C-SCORE complex, where the D1 domain of C-SCORE preferentially interacts with syntaxin (also see **Fig. 2*A***, where D1 spontaneously assembles onto syntaxin) with minimal interactions with the adjacent D2 domain, thus yielding a minimal increase in the FRET ratio. The stoichiometry of this complex could well be 2:2, owing to the high concentration of syntaxin (∼10 µM), which favors its oligomerization (18) (**Fig. 6*B***). We propose that Munc18 prevents the formation of these complexes, by acting as a chaperone that controls folding, and perhaps by inhibiting the oligomerization of syntaxin (**Fig. 6*A*** and ***B***), such that a robust increase in the FRET ratio was detected only when all four SNARE motifs were present (**Fig. 6*E***). These experiments highlight a unique role for Munc18 in orchestrating SNARE complex assembly.

### Munc18 promotes the assembly of D1 and D2 fragments of SNAP-25B into functional SNARE complexes

To address whether our *in vitro* folding results translate to functional outcomes, we turned to a well-established lipid mixing assay (49). We probed the impact of Munc18 on SNARE complex assembly by measuring lipid mixing between Syb- and syntaxin-bearing liposomes in the presence of the SNARE motifs of SNAP-25B. Neither individual SNARE motif of SNAP-25B, D1/D2, supported lipid mixing, either in the presence or absence of Munc18 (**Fig. 7*A*** and ***B***); when both D1 and D2 were present simultaneously, only a low level of lipid mixing occurred. Strikingly, under these same conditions, the addition of Munc18 dramatically enhanced the lipid mixing signal, as shown in **Fig. 7*C***. These results highlight the involvement of both SNARE motifs of SNAP-25B in the fusion reaction, perhaps via the action of Munc18 on syntaxin.

**Fig. 7.**
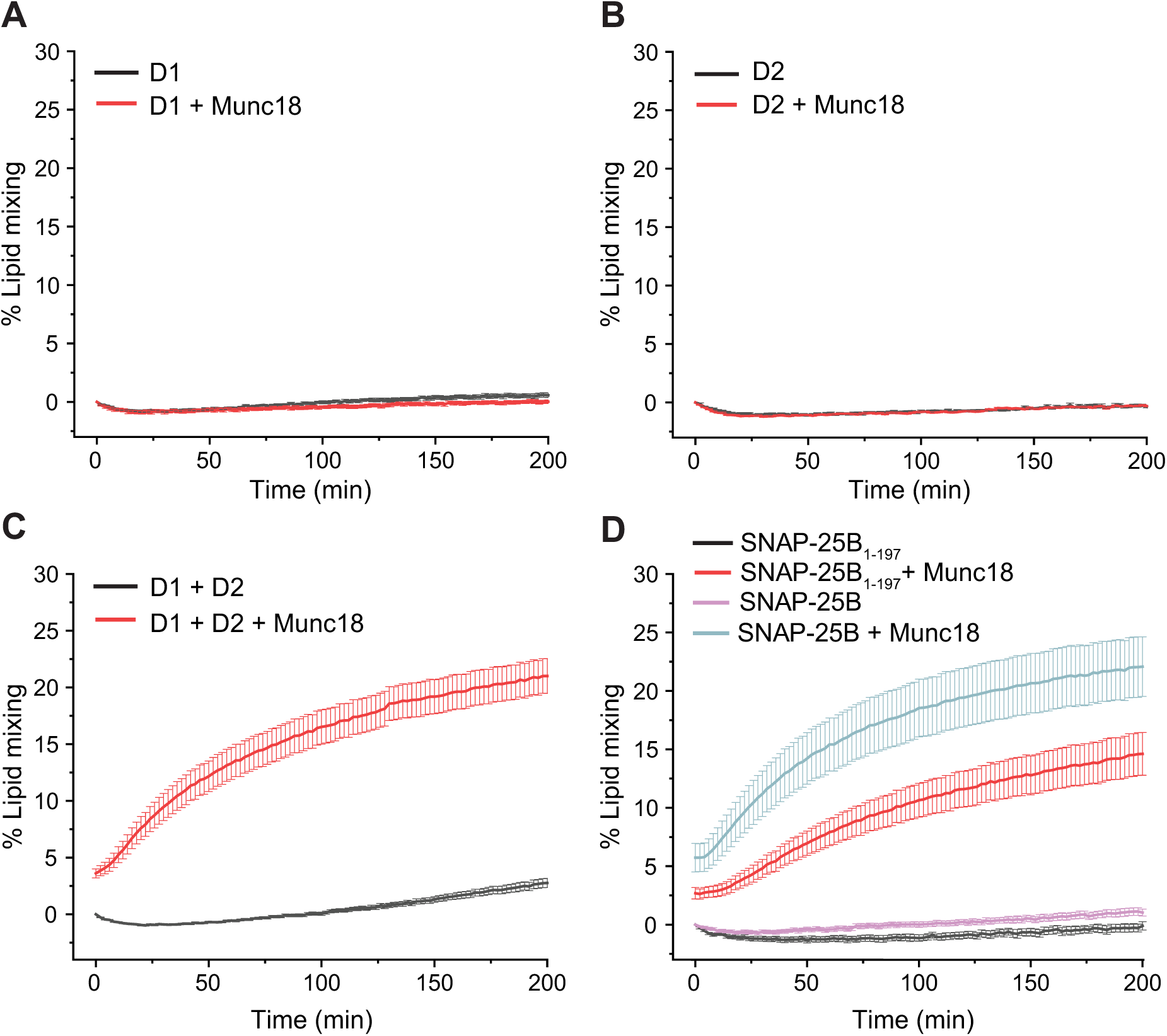
Munc18 promotes the assembly of full-length SNAP-25B, and fragments, into functional SNARE complexes. All experiments were conducted in the presence and absence of Munc18. In the minus Munc18 condition, v-SNARE and syntaxin-bearing liposomes were mixed in the presence of: (*A*) D1 (n = 12, N = 4), (*B*) D2 (n = 12, N = 4), (*C*) D1 + D2 (n = 27, N = 10), (*D*) full-length SNAP-25B (n = 10, N = 4) and SNAP-25B_1-197_ (n = 12, N = 4). In the plus Munc18 condition, the number of trails were: (*A*) D1 (n = 11, N = 4), (*B*) D2 (n = 12, N = 4), (*C*) D1 + D2 (n = 20, N = 7), (*D*) full-length SNAP-25B (n = 18, N = 6), and SNAP-25B_1-197_ (n = 18, N = 5). Error bars represent the standard error of the mean. n, total number of trials; N, number of sets of independent materials used to conduct the trials. In most cases, the same number of trials, 3, were conducted with each independent set of materials.

We further explored the structural elements of SNAP-25B that regulate SNARE assembly by focusing on the cleavage product of BoNT/A, SNAP-25B_1-197_; this truncated protein was analyzed in the standard lipid mixing assay (**Fig. 7*D***). We emphasize that SNAP-25B must be pre-complexed with syntaxin in order to drive membrane fusion in reconstituted fusion assays; the acute addition of soluble SNAP-25B to v- and syntaxin-bearing liposomes fails to result in fusion (26, 49, 50). As expected, and analogous to the WT protein, soluble SNAP-25B_1-197_ largely failed to support lipid mixing (26). Remarkably, in the presence of Munc18, we observed that soluble WT SNAP-25B efficiently assembled into functional SNARE complexes as evidenced by the robust lipid mixing signals. Moreover, SNAP-25B_1-197_ was also able to support some degree of lipid mixing, as compared to the WT control, but - again - only in the presence of Munc18. These results further corroborate the ability of Munc18 to act as a molecular chaperone during functional SNARE assembly; Munc18 help folds SNAP-25B, as well as the D1/D2 fragments (not-preassembled), into functional SNARE complexes that gave rise to robust lipid mixing signals. This positive role in congruent with the original observation that the yeast ortholog, Sec1 was required for fusion in yeast (51, 52).

## Discussion

Despite their central role in membrane fusion and synaptic transmission, the precise mechanism(s) by which neuronal SNAREs assemble into complexes remains unresolved. In the current study, we first examined the intrinsic pathway by which SNAREs alone assemble into productive complexes. We then addressed the question of how assembly is affected by the key regulatory protein, Munc18, which is absolutely required for membrane fusion *in vivo* (53). We began by conducting sequential order-of-addition experiments, using cytoplasmic domains and isolated SNARE motifs of the three major neuronal SNARE proteins, while monitoring interactions in real time. In the absence of regulatory proteins, these experiments revealed a pathway in which syntaxin and SNAP-25B first bind to each other and then cooperate to bind Syb (**Fig. 1*B-D***). However, this process became more intricate when we conducted assembly experiments using the isolated SNARE motifs of SNAP-25B; D1, and D2. We asked whether these motifs enter SNARE complexes at different time points to potentially introduce additional layers of regulation, or whether they join simultaneously. We found that labeled Syb folds more rapidly into the syntaxin-D1 complex versus the syntaxin-D2 complex, when syntaxin is added first, and D1 or D2 is added second, demonstrating that the two SNARE motifs of SNAP-25B are kinetically non-equivalent (**Fig. 1*E-G***) (18, 21). Interestingly, when D1 was labeled, syntaxin-D1 and H3-D1 formed dead-end complexes that failed to bind Syb and D2 (**Fig. 2*A*** and ***D***) (**Fig. 3**) (14). These experiments revealed that the order of addition, and differences in the concentrations of the reactants, can drastically affect the outcome in these assembly assays. These seemingly contradictory findings can be explained by an earlier structural study showing that H3•D1 can form four-helix bundles with a stoichiometry of 2:2 (13, 16) that would not be permissive for the incorporation of Syb (although D2 can enter on extremely long time scales). So, when low concentrations of labeled D1 are saturated with syntaxin or H3, non-productive complexes are formed, but when low concentrations of labeled Syb are first incubated with syntaxin, followed by the addition of D1, robust binding occurs. These order-of-addition experiments clearly revealed multiple parallel pathways by which SNAREs can assemble. A further complicating factor is the observation that at relatively high concentrations, syntaxin self-associates and forms a range of oligomers that appear to influence its interactions with other SNARE proteins. Because protein concentration can affect the outcome, we fixed these values in our experiments unless otherwise indicated.

The lack of affinity between syntaxin and D2 indicates that D2 joins the SNARE complex only after D1 has assembled in the context of full length SNAP-25B. Interestingly, D2 has little or no alpha helical structure, while D1 contains alpha helical structure near its N-terminus; perhaps this difference underlies the preferential and strong binding of D1 (18, 20, 21). Regardless, these findings result in a more nuanced model for the intrinsic assembly pathway: syntaxin first binds the N-terminal SNARE motif of SNAP-25B, followed by the binding of Syb and the C-terminal SNARE motif of SNAP-25B.

Interestingly, extending the D1 domain of SNAP-25B to residue 180 was sufficient to largely prevent, under these conditions, dead-end H3-D1 off-pathway interactions (**Fig. 4*B*** and ***D***). Since D1 enters the complex before D2 (**Fig. 1*E***) (18), these results indicate the formation of a nucleation site for Syb binding by the N-terminal region of the D2 domain (residues 141-180) in the H3-SNAP-25B_1-180_ complexes, providing evidence for directional N- to C-terminal zippering of v-SNARE•t-SNARE pairs. The key finding here is that the N-terminal region of D2 (within residues 141-180) acts as a gatekeeper to control the robust entry of Syb into the H3-D1 complexes. So, again, without any neighboring D2 sequences, D1 efficiently drives syntaxin (at either high or low concentrations) into the 2:2 off-pathway complex that is refractory to Syb binding. We emphasize that changing the order of addition can affect this outcome; starting with labeled Syb, then adding syntaxin and D1, did allow for ternary complex formation, likely along with other products that escaped detection.

We then expanded our experiments to address the role of the regulatory Habc domain of syntaxin, and the impact of Munc18 and Munc13-1_MUN_, on SNARE complex formation. Habc was originally thought to act as an autoinhibitory domain (38) that binds the H3 domain, resulting in the closed conformation of syntaxin that is refractory to SNARE complex assembly (24). Munc18 and a fragment of Munc13-1 were proposed to act together as chaperones that shift the equilibrium towards the open, activated form, of syntaxin (34, 41). We hypothesized that these regulatory proteins/domains might inhibit the formation of the off-pathway H3-D1 complex and thus favor the productive pathway. In the absence of Habc, neither Munc13_MUN_ nor Munc18 alone prevented the formation of off-pathway H3-D1 complexes (**Fig S6*C-F***). Importantly, in the presence of Habc, Munc13_MUN,_ and Munc18, isolated D1 did not interact with the isolated H3 domain, in the absence of other SNARE motifs, revealing the suppression of the H3-D1 off-pathway (**Fig. 5*A*** and ***C***). When the entire cytoplasmic domain of syntaxin was used, which includes the Habc domain, Munc18 alone prevented the formation of syntaxin-D1 off-pathway complexes (**Fig. 2*C***), and SNARE complex assembly was observed only when D1, D2, and Syb were concurrently present. Thus, inclusion of Munc18 dramatically changed the sequence of events leading to SNARE complex assembly, from a sequential to a concerted pathway, while the MUN domain of Munc13-1 was without effect in our experiments. We conclude that Munc18-mediated regulation of SNARE complex assembly primarily occurs via its interaction with the Habc domain of syntaxin. Notably, when the Habc and H3 domains of syntaxin are not linked, the assembly pathway is no longer concerted in the presence of Munc18 (**Fig. 5*B***, and ***D***). These findings, together with the complex behavior of D1 and D2, provide further insights into the modular nature of SNARE proteins, whereby only three components have multiple motifs and domains that give rise to complex assembly schemes.

To study the assembly of D1 and D2 in the context of the intact protein, we created C-SCORE, a SNAP-25B-based fluorescent sensor (**Fig. 6*C***) that reports SNAP-25B assembly into complexes based on the proximity between the D1 and D2 domains within the intact protein. As shown in **Fig. 6*D-E***, in the presence of Munc18, Syb and syntaxin must be present for the two SNARE motifs of SNAP-25B to robustly enter SNARE complexes. These experiments further support the concurrent SNARE assembly model dictated by the presence of Munc18 and Habc (**Fig. 2*C***).

Munc18 has been reported to play both positive and negative roles during SNARE complex formation. In the original yeast study, Sec1/Munc18 clearly played a positive role in exocytosis (26, 51, 52, 54). However, in subsequent studies based on murine proteins, Munc18 was shown to favor the closed conformation of syntaxin, a state that precludes interactions with other SNAREs, and it was argued that Munc18 functioned as a negative regulator of membrane fusion (24, 42, 55). Now, it has become apparent that Munc18 has several distinct modes of binding to syntaxin (24, 42, 43, 56), and the findings reported here and in ref. (26) reveal a net-positive role for Munc18 in SNARE assembly and lipid mixing, using reconstituted systems. Importantly, we found that the positive action of Munc18 relies on the Habc domain of syntaxin. Early studies of this domain suggested it impeded SNARE complex assembly (38), but more recent results revealed that Habc is essential for evoked exocytosis in neurons (57). Our findings provide a potential molecular explanation for this latter observation: Habc is required for Munc18 to activate syntaxin, consistent with an earlier model in which Munc13 and Munc18 serve to ‘open’ syntaxin, but here, only Munc18 was required, as we saw no effect of Munc13_MUN_. In addition, syntaxin is known to form clusters in the plasma membrane (58), potentially via liquid-liquid phase separation (LLPS) via the H3 domain (45); interestingly, this phase separation was inhibited by Munc18. Consistent with these findings, using FCS, we observed that Munc18 reduces the oligomerization of syntaxin (**Fig. 6*A*** and ***B***), and this de-clustering activity might also be involved in the activation of this protein. De-clustered syntaxin likely has Munc18 bound to it, perhaps via the Habc domain, to maintain it in an activated form.

Finally, we tested the functional implications of the Munc18-Habc-dependent concerted assembly model. Again, neither isolated D1 nor isolated D2 supported liposome fusion (**Fig. 7*A*** and ***B***), and the concurrent presence of D1 and D2 yielded a small but measurable lipid mixing signal. Strikingly, the addition of Munc18 greatly stimulated lipid mixing in the presence of D1 plus D2 (**Fig. 7*C***), and even with soluble SNAP-25B_1-197_ (**Fig. 7*D***), consistent with Munc18-driven assembly of SNARE complex. These findings also suggest that Munc18 can help SNAREs overcome the energy barrier for membrane fusion, to some degree, when the C-terminal of SNAP-25B is removed by BoNT/A. This observation helps to explain why BoNT/A treatment reduces but does not eliminate exocytosis (59).

We emphasize that the work reported here differs from an earlier landmark study that showed, for the first time, a positive effect of Munc18 on *in vitro* membrane fusion reactions when using preformed t-SNARE heterodimers (26). As explained earlier, preformed heterodimers are utilized because soluble SNAP-25B fails to support robust fusion in reconstituted lipid mixing assays (50), unless regulatory factors that drive SNARE complex assembly are added. To date, only two such factors have been described: synaptotagmin 1 (50), and – as shown here - Munc18 (**Fig. 7*D***). In the current study, the D1 and D2 domains, as well as full-length and truncated forms of SNAP-25B, were all added as soluble factors and were not pre-complexed with syntaxin. Therefore, Munc18 can stimulate lipid mixing by driving the folding of soluble SNAP-25B, as well as soluble D1 and D2 domains, into functional SNARE complexes formed via membrane-embedded syntaxin and Syb. These functional data are consistent with our emerging model: that in the presence of Munc18, all four SNARE motifs concurrently assemble into complexes to drive fusion. Hence, Munc18 appears to act as a chaperone, directly impacting the folding of SNARE proteins, mainly by interacting with syntaxin and enabling soluble SNAP-25B to be integrated into the fusion complex. (**Fig. 6*A*** and ***B***). For clarity, sequential and concerted assembly pathways are summarized in **Fig S8**. These pathways might become even more complex as additional regulatory factors are examined. Moreover, in cells, a variety of assembly pathways might occur.

We note that even in the presence of Munc18, SNARE complexes might be partially assembled during N- to C- terminal zippering. One interesting possibility is that the N-terminal region of D2 is partially assembled into SNARE complexes to potentially dock and prime the fusion machinery (60) and its full assembly is triggered by Ca^2+^. In this model, N-terminal assembly is concerted, while C-terminal assembly would be regulated, potentially by synaptotagmin 1, the major Ca^2+^ sensor for neuronal exocytosis (50, 61). More explicitly: in the absence of Ca^2+^, synaptotagmin 1 clamps fusion, potentially by hindering the full assembly of D2; then, in response to Ca^2+^, it might function to fold D2 into the SNARE complex (62). This model is supported by reconstitution studies, alluded-to above, showing that Ca^2+^•synaptotagmin facilitates the stable folding of SNAP-25B onto membrane-embedded syntaxin to drive fusion (50, 63). Munc18 is now the second protein identified that can directly regulate the folding of SNARE complexes (**Fig. 2*C*** and **Fig. 6*D***).

In future experiments, it will be important to include other key proteins, including synaptotagmin 1, in SNARE complex assembly assays. Additional factors like the ATPase NSF and its adaptor α-SNAP may also be involved in preventing the accumulation of off-pathway complexes by driving their ATP-dependent disassembly (64, 65). Finally, complexin preferentially binds to at least partially assembled SNARE complexes and might also control a late step in zippering (62).

## Methods

### Recombinant protein expression, purification and fluorescent labeling

cDNA constructs encoding cytoplasmic fragments of syntaxin-1A: residues 1–265 (syntaxin), 180-262 (H3), 1-180 (Habc); full-length SNAP-25B (1–206) and its fragments: residues 1-103 (D1), 1-137 (D1-Linker), 1-180 (SNAP-25B_1-180_), 1-197 (SNAP-25B_1-197_), 137-206 (D2); cytoplasmic domain of synaptobrevin 2: 1-96 (Syb); Munc18-1 (Munc18), and MUN domain of Munc13-1 (Munc13-1_MUN_, residues 859-1516 with 1408-1452 replaced by the sequence Glu-Phe (35, 66)) were cloned into a pET28 or pGEX-4T vector. The N-termini of each protein was fused with a His6-SUMO or GST tag. All proteins were expressed in *E. coli* BL21(DE3). Bacterial suspensions were grown in LB media at 37 °C until the OD_600_ reached 0.6; 500 µM isopropyl β-D-1-thiogalactopyranoside (IPTG) (GoldBio, I2481C) was then added, and proteins were expressed for 16 h at 20 °C. Bacterial pellets were collected by centrifugation, and sonicated (90 sec, 50% duty cycle) in resuspension buffer comprising 25 mM HEPES buffer at pH 7.4, 400 mM KCl, 5% glycerol, a protease inhibitor cocktail (PIC) (Roche, 046693132001), DNase, and RNase (15 µg/ml each). One % (v/v, final) Triton X-100 (Thermo Fisher Scientific, A16046) was mixed with the bacterial lysates, and the suspensions were rotated for 2 hours at 4 °C. The extracts were centrifuged at 31,000 × *g* for 45 min at 4 °C, and the supernatants were mixed with TALON Metal affinity resin (Takara, 635504) or Glutathione Sepharose^TM^ 4B (Cytiva, 17075604) at 4 °C for 2 h. The beads were washed with wash buffer (25 mM HEPES, pH 7.4, 400 mM KCl, 5% glycerol); 5 mM imidazole was added to the wash buffer for His-SUMO fused proteins. Finally, beads were treated with 0.5 µM recombinant SUMO protease (senp2) for His-SUMO tagged proteins, or 10 units/ml of thrombin for GST fusion proteins, at 4 °C overnight to cleave the protein of interest from the fusion tag (SUMO or GST). The eluted proteins were dialyzed against 25 mM HEPES, pH 7.4, 100 mM KCl, subjected to SDS-PAGE, and stained with Coomassie blue using bovine serum albumin (BSA) as a concentration standard.

To generate the C-SCORE construct of SNAP-25B, a 12 amino acid (-GSS-) flexible linker was inserted right after residue 84, followed by cyan fluorescent protein (CFP). A 20 amino acid (- GSS-) flexible linker was added at the C-terminus of CFP, followed by residues 85-206. The C-terminus of the modified SNAP-25B was then fused to Venus via a 12 amino acid (-GSS-) flexible linker. C-SCORE was prepared as an N-terminally tagged GST fusion protein and purified via the protocol described above.

Proteins were labeled (*) with Alexa Fluor 488 C5 maleimide at a Cys residue placed in SNARE domains as follows: D1 (M7C); D2 (R136C); cytoplasmic domain of Syb: S28C or T79C; and on an additional Cys residue placed at the C-terminal of the SNARE motif of syntaxin (H3, 180-262). Twenty µM of each protein was mixed with ∼240 µM of the Alexa Fluor 488 dye at 4 °C overnight. Free dye was removed using a PD-10 desalting column (Cytiva) followed by dialysis using an appropriate molecular weight cut-off membrane for 48 hours in 25 mM HEPES, pH 7.4, 100 mM KCl.

### Fluorescence anisotropy

Fluorescence anisotropy was used to monitor real-time interactions between SNAREs and regulatory proteins. Fluorescently labeled samples were excited at 488 nm, and the emission was monitored at 520 nm using a Fluorolog-QM (Horiba)l; all slits were 5 nm. The fluorescence anisotropy was calculated using the equation shown below, where ‘I’ is the fluorescence intensity, and H and V indicate horizontally and vertically polarized light, respectively. The first letter (subscript) defines the direction of excitation light, while the second letter indicates the direction of light detection. The anisotropy values were corrected for the instrument correction factor (G-factor, G), which was calculated as 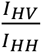

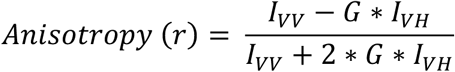

Following each addition, the fluorescence anisotropy was monitored in real time until a plateau was reached to obtain steady-state values. From each trace, we calculated the mean plateau anisotropy from the last 10 minutes of the measurement. Anisotropy changes were calculated by subtracting this mean value from the average anisotropy value of the corresponding isolated, labeled SNARE motif. In some cases, the rate of change was also determined, as described in the text. Each figure shows a representative trace and the number of independent trials; statistical analyses are provided in each of the Figure legends. In all the experiments, concentration of labeled protein was maintained between 200-300 nM while unlabeled proteins were used at ∼ 10 μM, unless otherwise indicated.

### *In vitro* FRET measurement

All FRET measurements were performed in a Fluorolog-QM (Horiba) equipped with DUAL monochromators. We measured the emission spectra of the sample (e.g. C-SCORE ± Munc18, C-SCORE ± Munc18 + syntaxin, and C-SCORE ± Munc18 + syntaxin + Syb) with a 1s integration time; slits were set to 5 nm. We calculated the ratio of the fluorescence intensity at 527 nm to 477 nm and used it as a measure of interaction between the D1 and D2 domain of C-SCORE during SNARE complex formation.

### Measurement of diffusion time by fluorescence correlation spectroscopy (FCS)

The diffusion time of labeled H3 was measured using an FCS instrument built in-house, as described (44). Briefly, we use a polarized continuous wave 488 nm laser beam to excite fluorescently labeled samples via a 1.2 NA objective lens (Olympus, USA). Fluorescence signals were collected by the two co-aligned objective lenses and acquired in two channels using single photon avalanche photodiode detectors (APD, PerkinElmer Inc.). The fluorescence intensity time trace was recorded

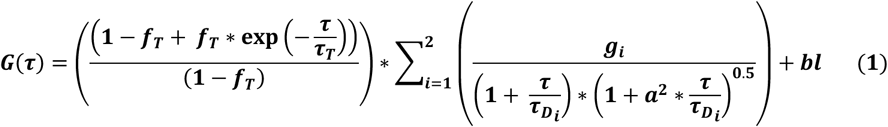

using a hardware correlator (PicoHarp 300), and the microsecond correlation function was calculated using SymPhoTime64 software (PicoQuant, Germany). We analyzed the FCS data using equation 1, where *G*(*τ*) represents the autocorrelation function and *τ* is the lag time. The analysis was performed using Origin Pro software (Northampton, MA, USA) to measure the following parameters; *f_T_* : triplet-state fraction; *τ_T_* : triplet-state relaxation time, *τ_D_* : diffusion time of the fluorescent molecule, and *g*: amplitude of diffusion component. In the equation, ‘a’ serves as the structure factor and ‘bl’ as the background.

For FCS experiments, a stock solution of Alexa Fluor 488 labeled H3 (∼ 10 μM) was diluted to ∼ 50 nM in HEPES buffer (25 mM HEPES, 100 mM KCl). The concentrations of unlabeled proteins, Habc or Munc18, were kept at ∼ 10 μM. Each FCS trace was recorded for five minutes and analyzed using the above equation considering two diffusing species (*i* = 2). Fractional population of each species, e.g. *g_2_*, was calculated as *g_2_*/(*g_1_*+*g_2_*).

### Preparation of SNARE-bearing proteoliposomes

All lipids were purchased from Avanti Polar Lipids. v-SNARE bearing FRET donor/acceptor liposomes were prepared using a lipid mixture comprising DOPC (1,2-dioleoyl-sn-glycero-3-phosphocholine), DOPS (1,2-dioleoyl-sn-glycero-3-phospho-L-serine), Rhodamine-PE (1,2-dioleoyl-sn-glycero-3-phosphoethanolamine-N-(lissamine rhodamine B sulfonyl) (ammonium salt), and NBD-PE (1,2-dioleoyl-sn-glycero-3-phosphoethanolamine-N-(7-nitro-2-1,3-benzoxadiazol-4-yl)) at a molar ratio of 77:20:1.5:1.5. Chloroform stocks of lipids were mixed at the appropriate ratio and dried under nitrogen. Lipid films were then dried in a lyophilizer for two hours to remove any residual chloroform or moisture. The dried lipid film was resuspended in detergent (1% w/v, octyl-β-D-glucopyranoside (OG), CHEM IMPEX) containing full length Syb at a protein:lipid ratio of 1:500. The mixture was diluted 3-fold using reconstitution buffer (25 mM HEPES and 100 mM KCl) to reduce the detergent concentration below its CMC (0.53%, w/v), and the detergent was removed via extensive dialysis against reconstitution buffer. Proteoliposomes were then separated from free protein via flotation on a step gradient of 40%, 30% and 0% Accudenz (Accurate Chemical & Scientific corporation); the top layer (0% Accudenz), containing the proteoliposomes, was collected. Syntaxin-bearing proteoliposomes were generated in the same manner, but with DOPC and DOPS at a molar ratio of 80:20.

### *In vitro* lipid mixing assays

Each fusion reaction consisted of 2 μl of purified Syb proteoliposomes and 50 μl of syntaxin proteoliposomes, along with accessary proteins and soluble SNAP-25B or SNAP-25B fragments, in a final volume of 200 μl, in black bottom 96 well plates. All components of the reaction were preincubated at 4 °C for 3 hours, and then fusion reactions were initiated by inserting the plate in a plate reader set to 37°C. Lipid mixing was monitored by following dequenching of the NBD (λ_em_ = 538 nm) due to loss of FRET during fusion. One % n-dodecylmaltoside (w/v) was added at the end of each assay to completely dequench the NBD fluorescence, and this value was used to calculate the lipid mixing percentage.

### Statistical analysis

The number of independent measurements and statistical analyses are provided in Figure legends. Statistical analyses were performed using Excel and GraphPad Prism.

## Acknowledgments

We thank Dr. Nikunj Mehta for designing the labeling sites in SNARE proteins and helping with cDNA constructs for protein expression. We thank all members of the Chapman lab for valuable discussions and feedback regarding this manuscript. This study was supported by grants from the NIH (MH061876 and NS136306) to E.R.C. E.R.C. is an Investigator of the Howard Hughes Medical Institute.

## Author Contributions

**Vicky Vishvakarma:** Conceptualization; Investigation; Writing original draft; Writing and editing.

**Edwin R. Chapman:** Conceptualization; Funding acquisition; Supervision; Resources; Writing original draft; Project administration; Writing and editing.

