## Supplementary Information for "Munc18 reprograms the intrinsic neuronal SNARE complex assembly pathway"

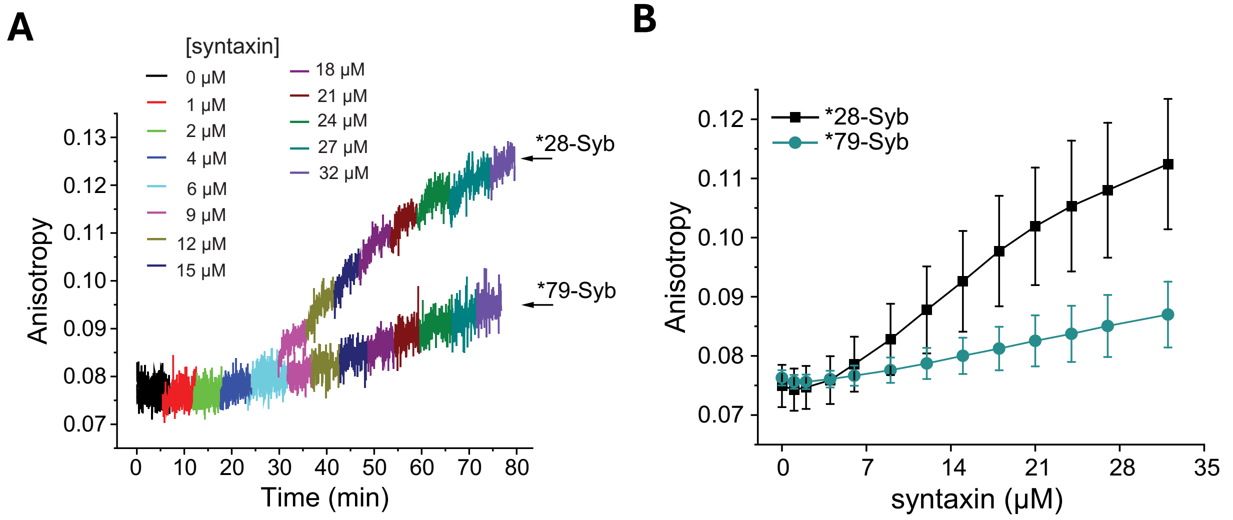

**Fig. S1.** (*A*) Syntaxin-Syb interactions monitored as a function of time at different [syntaxin] via fluorescence anisotropy of Syb, labeled (*) at either amino acid position 28 (n = 3) or 79 (n = 4) with Alexa Fluor 488. The [syntaxin] is indicated in the color key. (*B*) Analysis of the data represented in panel *A*. Error bars represent standard deviation.

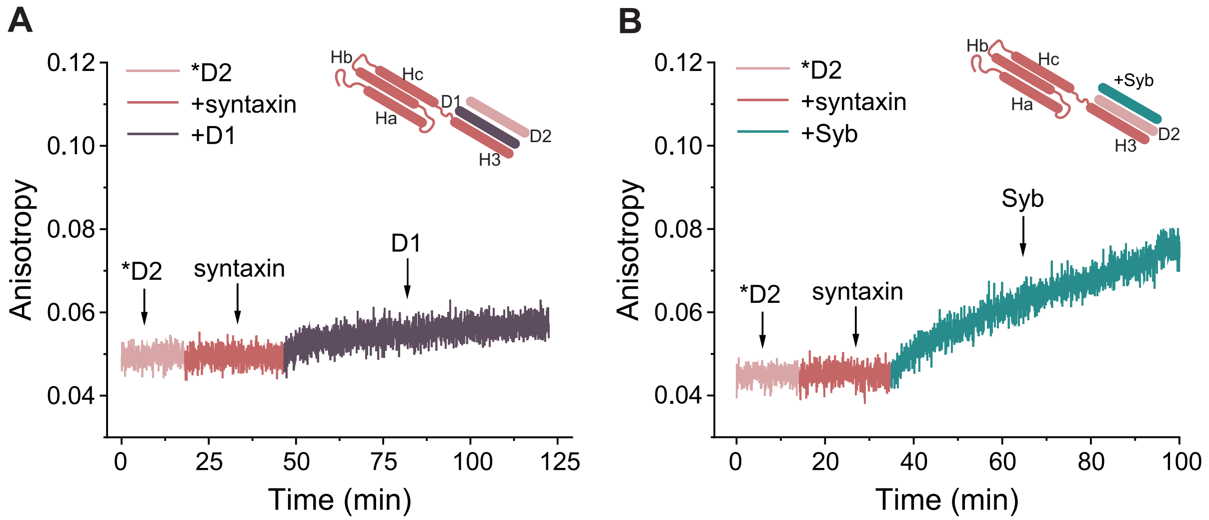

**Fig. S2.** The D2 domain of SNAP-25B exhibits ternary interactions with syntaxin-D1 and syntaxin-Syb, with no binary interaction with syntaxin. (*A*) Fluorescence anisotropy of labeled (*) D2 domain measured with sequential addition of syntaxin and D1; n = 6. (*B*) Fluorescence anisotropy of labeled D2 domain measured with sequential addition of syntaxin and Syb; n = 3. These data are included in the plot shown in **Fig. 2*D***.

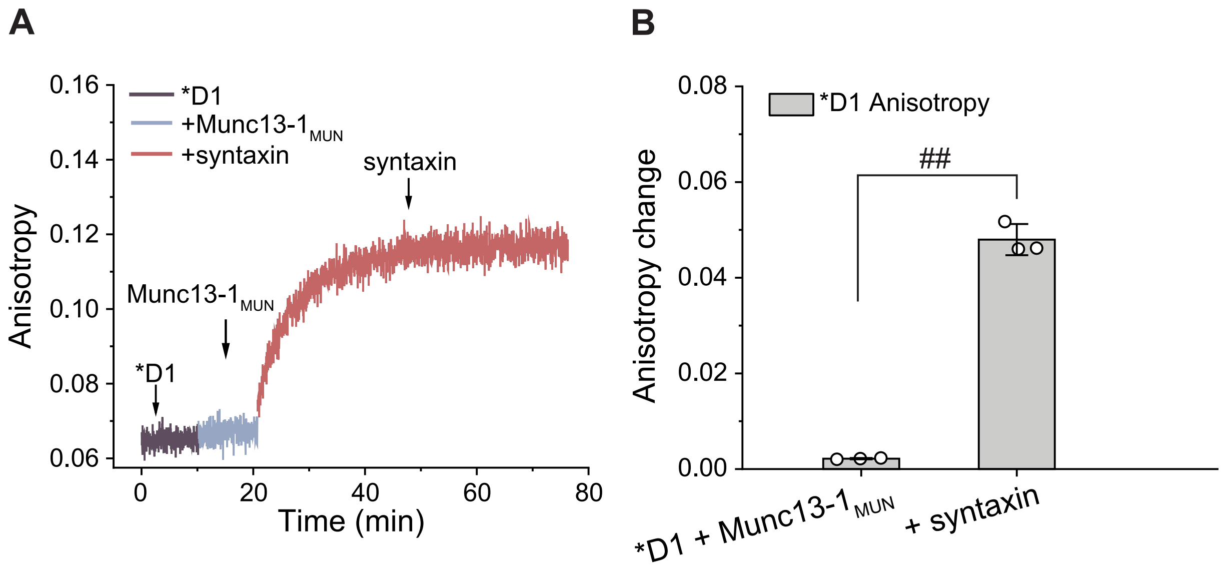

**Fig. S3.** Munc13-1$_{MUN}$ does not block syntaxin-D1 interactions. (*A*) Fluorescence anisotropy of labeled (*) D1 measured with the sequential addition of Munc13-1$_{MUN}$ and syntaxin; n = 3. (*B*) Quantification of the change in fluorescence anisotropy of D1 corresponding to the experiment shown in panel (*A*). Error bars represent the standard deviation. Statistical significance was determined by Welch's unpaired t-test; $^{\#}p < 0.05$, $^{\#\#}p < 0.01$, $^{\#\#\#}p < 0.001$, $^{\#\#\#\#}p < 0.0001$, ns-not significant.

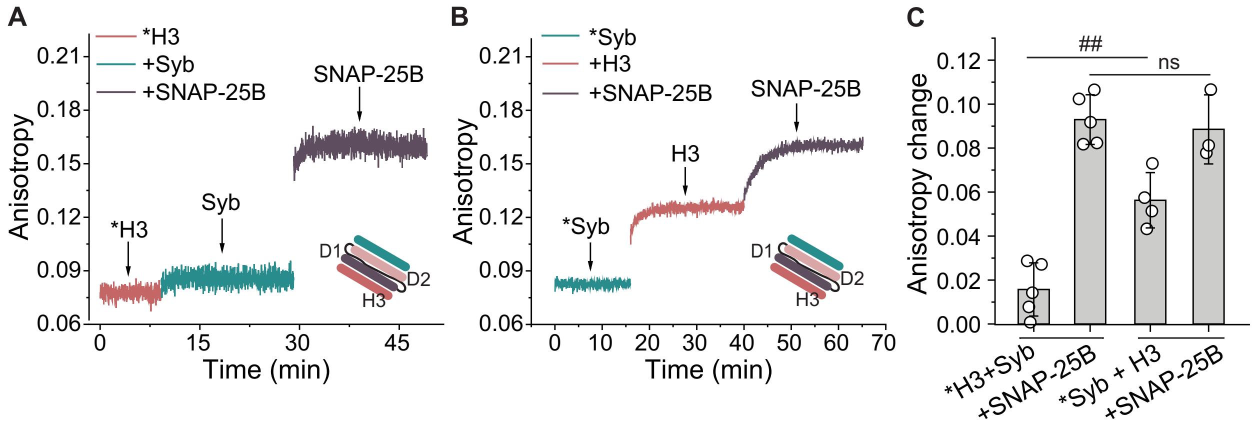

**Fig. S4**. The concentration of H3 affects how it binds to Syb. (*A*) Fluorescence anisotropy of labeled (*) H3 (~200 nM) measured with sequential addition of Syb (10 µM), and SNAP-25B (10 µM); n = 5. (*B*) Fluorescence anisotropy of labeled (*) Syb (~200 nM) measured with sequential addition of unlabeled H3 (10 µM), and SNAP-25B (10 µM); n = 3. Note: the anisotropy change for the Syb-H3 interaction in panel (*A*) is smaller than in panel (*B*), because in the latter panel the higher [H3] forms oligomers; it is also possible the oligomers bind more tightly to Syb. (*C*) Quantification of the fluorescence anisotropy changes of labeled H3 or Syb from panels (*A*) and (*B*). The line segments at the bottom indicate the grouping of the same order-of-addition experiments. Error bars represent the standard deviation. Statistical significance was determined by Welch's unpaired t-test; $^{\#}p < 0.05$, $^{\#\#}p < 0.01$, $^{\#\#\#}p < 0.001$, $^{\#\#\#\#}p < 0.0001$, ns-not significant.

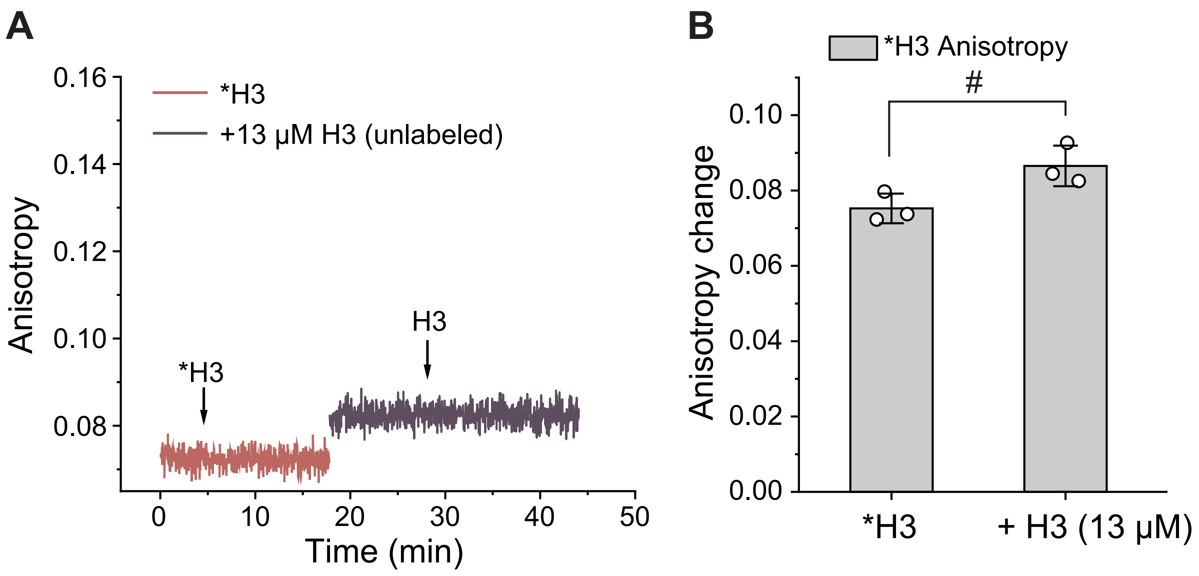

**Fig. S5.** The SNARE motif of syntaxin (H3) multimerizes. (*A*) Fluorescence anisotropy of labeled (*) H3 measured before and after the addition of 13 µM unlabeled H3; n = 3. (*B*) Change in fluorescence anisotropy of H3 after addition of 13 µM of unlabeled H3. The error bar represents the standard deviation. Statistical significance was determined by Welch's unpaired t-test; $^{\#}p < 0.05$, $^{\#\#}p < 0.01$, $^{\#\#\#}p < 0.001$, $^{\#\#\#\#}p < 0.0001$, ns-not significant.

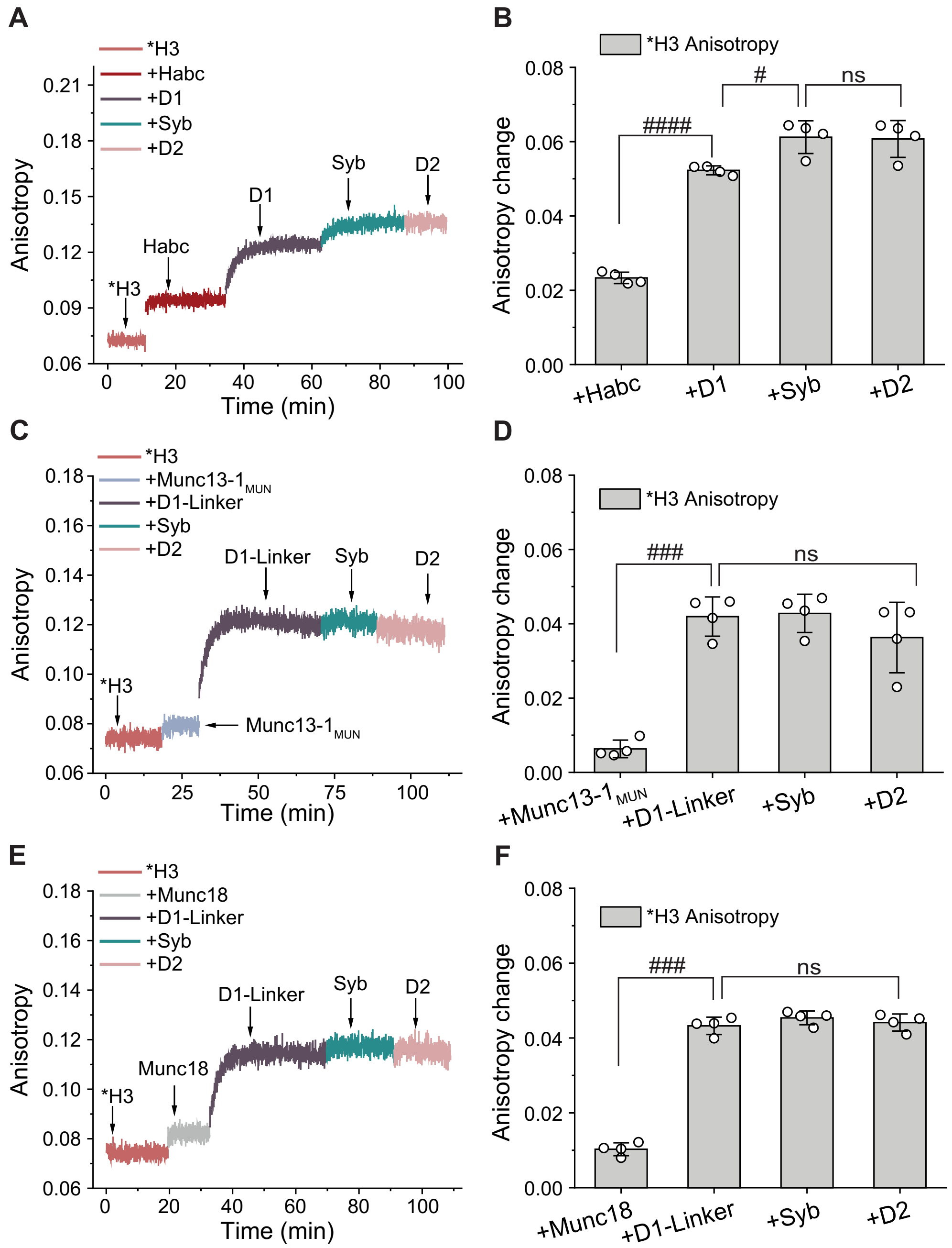

**Fig. S6.** The syntaxin Habc domain allows Syb to enter the H3-D1-linker complex; Munc13-$1_{\mathrm{MUN}}$ was without effect, and Munc18 failed to inhibit the H3-D1 off pathway in the absence of Habc. Left panels (*A, C, E)*: Fluorescence anisotropy traces of labeled (*) H3 measured with the sequential addition of: (*A*) the syntaxin Habc domain (n = 4), (*C*) the SNARE binding domain of Munc13-1 (MUN domain) (n = 4), and (*E*) Munc18 (n = 4); in each case these additions were followed by the sequential addition of D1/D1-Linker, Syb, and D2. Right panels (*B*, *D*, *F*): Quantification of the change in fluorescence anisotropy of H3 from the corresponding panels (*A*), (*C*), and (*E*), respectively. Error bars represent the standard deviation. Statistical significance was determined by Welch's unpaired t-test; $^{\#}p < 0.05$, $^{\#\#}p < 0.01$, $^{\#\#\#}p < 0.001$, $^{\#\#\#\#}p < 0.0001$, ns-not significant.

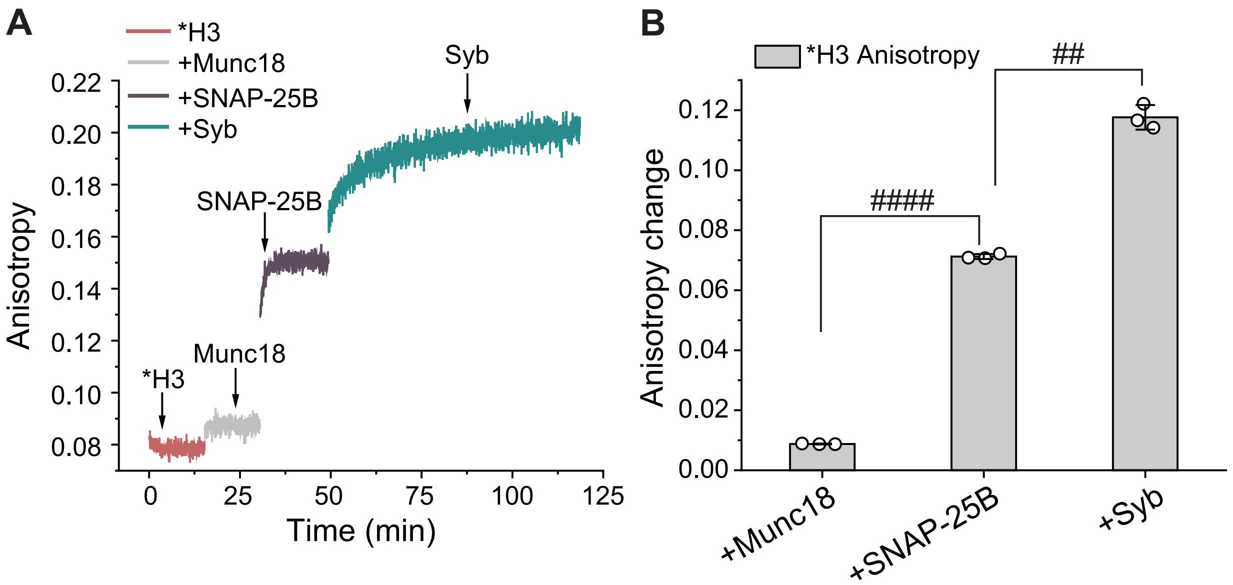

**Fig. S7.** Munc18 does not block H3-SNAP-25B interactions in the absence of the Habc domain of syntaxin. (*A*) Fluorescence anisotropy of labeled (*) H3 measured with the sequential addition of Munc18, SNAP-25B, and Syb; n = 3. (*B*) Quantification of the change in fluorescence anisotropy of H3 corresponding to the experiment shown in panel (*A*). Error bars represent the standard deviation. Statistical significance was determined by Welch's unpaired t-test; $^{\#}p < 0.05$, $^{\#\#}p < 0.01$, $^{\#\#\#}p < 0.001$, $^{\#\#\#\#}p < 0.0001$, ns-not significant.

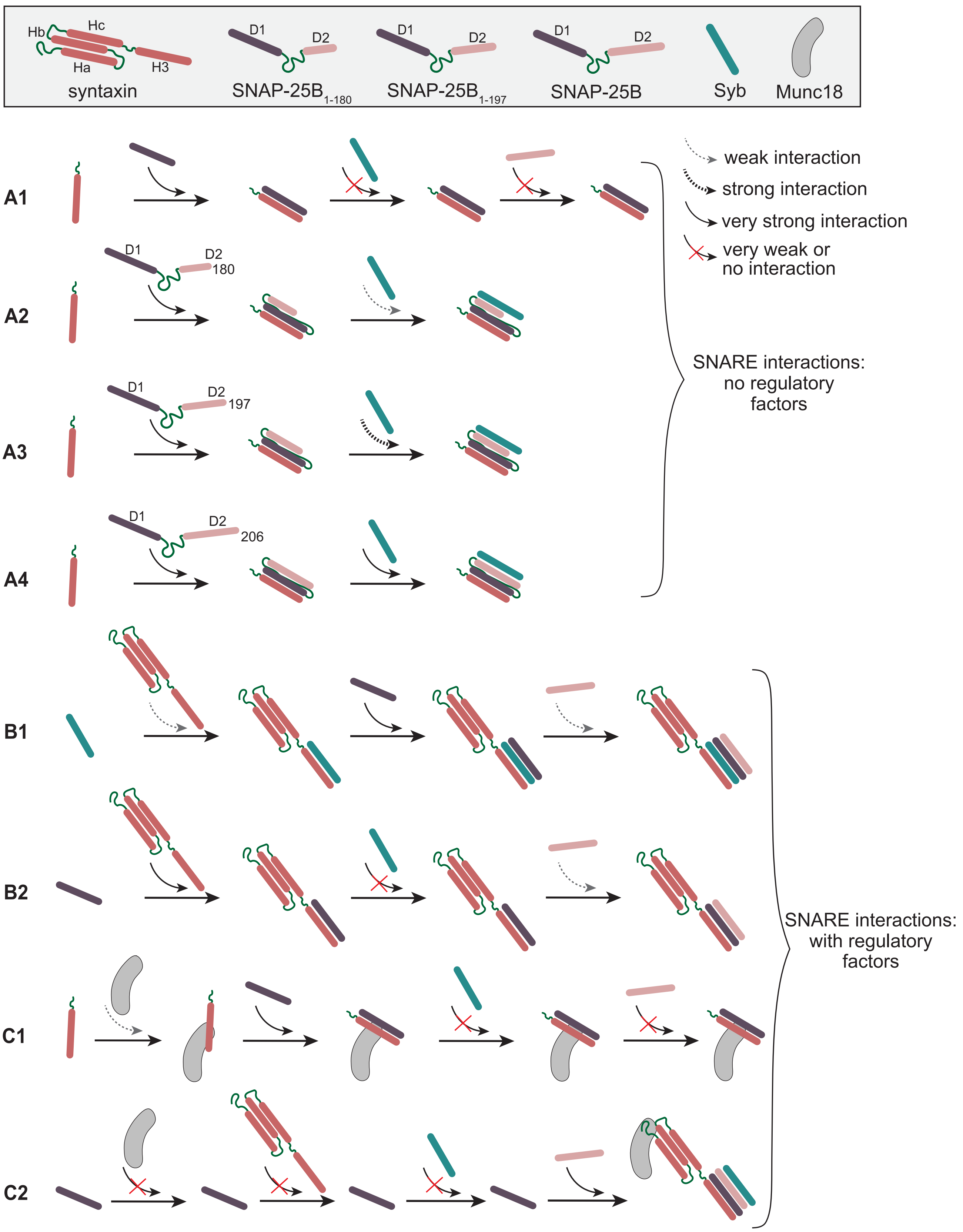
Hb
Hc
Ha
H3
syntaxin
D1
D2
SNAP-25B1-180
D1
D2
SNAP-25B1-197
D1
D2
SNAP-25B
Syb
Munc18
weak interaction
strong interaction
very strong interaction
very weak or no interaction
A1
A2
D1
D2
180
A3
D1
D2
197
A4
D1
D2
206
SNARE interactions: no regulatory factors
B1
B2
C1
C2
SNARE interactions: with regulatory factors

**Fig. S8.** Model describing interactions among SNARE proteins in the presence and absence of regulatory factors. The proteins used in this study are shown in the shaded box at the top. (*A1*-*A4*) H3-D1 forms an off-pathway complex, that is refractory for binding to D2 or Syb; it is shown as a simplified 1:1 complex but is likely to be the previously characterized 2:2 complex (14). Extension of D1 towards full-length SNAP-25B enhances the binding of Syb. (*B1*-*B2*) SNARE motif interactions in the presence of the Habc domain of syntaxin. The formation of off-pathway complexes depends on the order in which components were added. (*C1*-*C2*) Munc18 gates the entry of syntaxin into SNARE complexes by interacting with the Habc domain. The D1-syntaxin interaction is inhibited via Munc18-Habc interactions. In the absence of Habc, Munc18 does not inhibit formation of the H3-D1off-pathway complex.

**Table S1:** Diffusion parameters obtained from FCS data using equation 1 (see Methods for details)

| ***Sample Name*** | $g_1$ | $\tau_{D1}$ (μs) | $g_2$ | $\tau_{D2}$ (μs) | **Fractional population of large H3 oligomers ($g_2/(g_1+g_2)$)** |
|---|---|---|---|---|---|
| *H3 | 1.70 | 28.28 | 0.70 | 175.19 | |
| | 1.62 | 28.00 | 0.81 | 123.95 | |
| | 1.70 | 28.00 | 0.76 | 134.30 | |
| | 0.35 | 108.90 | 0.03 | 1612.59 | 0.0685 |
| | 0.30 | 75.94 | 0.11 | 266.46 | |
| | 0.41 | 104.45 | 0.03 | 1690.58 | 0.0593 |
| | 0.86 | 71.39 | 0.29 | 4142.31 | 0.2528 |
| | 0.93 | 53.95 | 0.08 | 1998.75 | 0.0834 |
| | 0.28 | 91.89 | 0.04 | 1529.40 | 0.1334 |
| | 0.26 | 100.51 | 0.03 | 2409.79 | 0.0890 |
| | 0.21 | 66.17 | 0.07 | 369.14 | |
| | 0.07 | 60.26 | 0.06 | 252.26 | |
| | 0.07 | 60.91 | 0.05 | 256.16 | |
| | 0.90 | 79.61 | 0.17 | 3540.51 | 0.1587 |
| | 0.12 | 119.66 | 0.01 | 2656.65 | 0.0673 |
| | 0.11 | 122.15 | 0.00 | 1405.98 | 0.0350 |
| | 0.12 | 118.74 | 0.01 | 1021.00 | 0.0608 |
| | 0.11 | 115.04 | 0.01 | 1103.19 | 0.0568 |
| | | | 0.32 | 132.92 | |
| | | | 0.28 | 127.55 | |
| | | | 0.25 | 132.19 | |
| | 0.08 | 50.00 | 0.14 | 170.59 | |
| | 0.11 | 65.99 | 0.09 | 221.61 | |
| | 0.10 | 68.43 | 0.07 | 225.25 | |
| | 0.18 | 109.82 | 0.02 | 2055.48 | 0.1187 |
| | 0.07 | 60.15 | 0.11 | 175.59 | |
| | 0.06 | 48.28 | 0.13 | 164.70 | |
| | 0.07 | 70.44 | 0.05 | 210.75 | |
| | 0.09 | 89.47 | 0.02 | 375.11 | |
| | 0.08 | 79.07 | 0.03 | 282.75 | |

| | | | | | |
|---|---|---|---|---|---|
| | $g_1$ | $\tau_{D1}$ (μs) | $g_2$ | $\tau_{D2}$ (μs) | Fractional population of large H3 oligomers ($g_2/(g_1+g_2)$) |
| *H3 + Munc18 | 0.18 | 117.50 | 0.01 | 19553.58 | 0.0317 |
| | 0.16 | 80.00 | 0.03 | 369.82 | |
| | 0.09 | 34.43 | 0.11 | 181.50 | |
| | 0.08 | 28.60 | 0.13 | 165.96 | |
| | 0.06 | 36.11 | 0.08 | 172.93 | |
| | 0.06 | 30.00 | 0.09 | 160.38 | |
| | 0.05 | 25.65 | 0.09 | 159.91 | |
| | 0.05 | 26.33 | 0.09 | 161.69 | |
| | 0.06 | 38.38 | 0.08 | 187.33 | |
| | 0.12 | 117.19 | 0.02 | 43011.04 | 0.1446 |
| | 0.06 | 36.00 | 0.08 | 172.74 | |
| | 0.23 | 150.96 | 0.01 | 43014.82 | 0.0500 |
| | 0.17 | 94.42 | 0.06 | 364.62 | |
| | 0.09 | 59.51 | 0.15 | 196.95 | |
| | 0.04 | 50.47 | 0.10 | 182.12 | |
| | 0.06 | 63.37 | 0.08 | 202.38 | |
| | 0.06 | 71.33 | 0.06 | 226.31 | |
| | 0.05 | 59.09 | 0.08 | 187.60 | |
| | 0.03 | 41.06 | 0.10 | 166.64 | |
| | 0.07 | 55.89 | 0.14 | 213.26 | |
| | 0.08 | 58.62 | 0.13 | 225.60 | |
| | 0.10 | 72.31 | 0.11 | 251.25 | |
| | $g_1$ | $\tau_{D1}$ (μs) | $g_2$ | $\tau_{D2}$ (μs) | Fractional population of large H3 oligomers ($g_2/(g_1+g_2)$) |
| *H3 + Habc | 0.08 | 140.38 | 0.03 | 4241.86 | 0.2584 |
| | 0.08 | 137.27 | 0.03 | 3730.91 | 0.2485 |
| | 0.08 | 132.20 | 0.03 | 3379.54 | 0.2416 |
| | 0.08 | 133.85 | 0.02 | 3243.21 | 0.2324 |
| | 0.08 | 146.67 | 0.02 | 5201.87 | 0.2248 |

| | | | | | |
|---|---|---|---|---|---|
| | 0.33 | 143.40 | 0.12 | 2788.85 | 0.2707 |
| | 0.34 | 135.67 | 0.12 | 2009.80 | 0.2625 |
| | 0.34 | 145.33 | 0.12 | 3120.36 | 0.2614 |
| | 0.34 | 132.11 | 0.13 | 2092.59 | 0.2747 |
| | 0.36 | 137.66 | 0.13 | 2591.32 | 0.2675 |
| | 0.38 | 157.91 | 0.13 | 4736.93 | 0.2520 |
| | 0.36 | 159.33 | 0.13 | 3776.96 | 0.2594 |
| | 0.35 | 155.42 | 0.14 | 4411.95 | 0.2792 |
| | 0.27 | 195.81 | 0.09 | 7370.68 | 0.2572 |
| | 0.29 | 165.40 | 0.09 | 3881.17 | 0.2325 |
| | 0.28 | 196.50 | 0.10 | 7689.85 | 0.2597 |
| | 0.28 | 204.09 | 0.12 | 9208.49 | 0.2967 |
| | 0.28 | 244.10 | 0.13 | 15914.67 | 0.3072 |
| | 0.30 | 152.79 | 0.09 | 2941.89 | 0.2393 |
| | 0.30 | 146.54 | 0.10 | 2809.81 | 0.2529 |
| | 0.30 | 147.52 | 0.10 | 3024.89 | 0.2512 |
| | 0.32 | 150.52 | 0.11 | 3085.93 | 0.2512 |
| | **$g_1$** | **$\tau_{D1}$ (μs)** | **$g_2$** | **$\tau_{D2}$ (μs)** | **Fractional population of large H3 oligomers ($g_2/(g_1+g_2)$)** |
| *H3 + Habc + Munc18 | 0.09 | 165.29 | 0.02 | 1106.46 | 0.1972 |
| | 0.08 | 144.77 | 0.04 | 632.39 | |
| | 0.08 | 142.56 | 0.04 | 683.26 | |
| | 0.08 | 146.17 | 0.03 | 693.69 | |
| | 0.01 | 96.67 | 0.02 | 322.72 | |
| | 0.01 | 83.20 | 0.03 | 315.32 | |
| | 0.07 | 87.75 | 0.12 | 334.23 | |
| | 0.07 | 94.66 | 0.12 | 342.72 | |
| | 0.06 | 81.62 | 0.13 | 326.21 | |
| | 0.06 | 73.48 | 0.14 | 307.57 | |
| | 0.06 | 77.62 | 0.13 | 321.69 | |
| | 0.04 | 90.25 | 0.09 | 275.05 | |
| | 0.02 | 59.32 | 0.11 | 252.21 | |
| | 0.03 | 79.21 | 0.10 | 260.90 | |
| | 0.04 | 81.60 | 0.10 | 261.24 | |
